# Large haploblocks underlie rapid adaptation in an invasive weed

**DOI:** 10.1101/2022.03.02.482376

**Authors:** Paul Battlay, Jonathan Wilson, Vanessa C. Bieker, Christopher Lee, Diana Prapas, Bent Petersen, Sam Craig, Lotte van Boheemen, Romain Scalone, Nissanka P. de Silva, Amit Sharma, Bojan Konstantinović, Kristin A. Nurkowski, Loren H. Rieseberg, Tim Connallon, Michael D. Martin, Kathryn A. Hodgins

## Abstract

Adaptation is the central feature and leading explanation for the evolutionary diversification of life. Adaptation is also notoriously difficult to study in nature, owing to its complexity and logistically prohibitive timescale. We leverage extensive contemporary and historical collections of *Ambrosia artemisiifolia*—an aggressively invasive weed and primary cause of pollen-induced hayfever—to track the phenotypic and genetic causes of recent local adaptation across its native and invasive ranges in North America and Europe, respectively. Large haploblocks— indicative of chromosomal inversions—contain a disproportionate share (26%) of genomic regions conferring parallel adaptation to local climates between ranges, are associated with rapidly adapting traits, and exhibit dramatic frequency shifts over space and time. These results highlight the importance of large-effect standing variants in rapid adaptation, which have been critical to *A. artemisiifolia*’s global spread across vast climatic gradients.

## INTRODUCTION

Adaptation can be rapid, yet changes in the traits and genes that underlie adaptation are difficult to observe in real time because speed is relative: that which is fast in evolutionary terms is slow from the human perspective. Thus, while we know that adaptation is central to evolutionary diversification, species persistence and invasiveness, the genetic and phenotypic dynamics of adaptation are difficult to document outside of the laboratory.

Invasive species are powerful systems for characterizing adaptation in nature, owing to several features that make them unique. In particular, biological invasions coincide with exceptionally rapid evolution^1–3^, observable over human lifespans, as invasive populations can encounter drastically different environmental conditions from those of their source populations. Many invasions are, moreover, documented in large, geo-referenced herbarium collections, which can be phenotyped and sequenced to identify and track adaptive evolutionary changes through time—feats that are rarely achieved in natural populations. Invasive species frequently inhabit broad and climatically diverse ranges, which favours the evolution of adaptations to local environmental conditions^2,3^, along with evolved tolerance of environmental extremes, which may be conducive to invasiveness^4^. Because they often occupy geographically broad native and invasive ranges, invasive species allow for tests of the predictability of evolution—a major puzzle in biology—as local adaptation across native and invasive ranges may favour either parallel or unique genetic solutions to shared environmental challenges. However, despite the promise of historical records and other features of invasive species that make them tractable systems for capturing adaptation in action, this treasure trove of data has not been fully utilized to elucidate the genetic basis of local adaptation during recent range expansions.

*Ambrosia artemisiifolia*, an annual weed native to North America, has mounted successful invasions on all continents except Antarctica^5^. The species thrives in disturbed habitats and has experienced extensive range shifts, historically documented in pollen records and herbarium collections. It also produces highly allergenic pollen, which is the chief cause of seasonal allergic rhinitis and asthma in the United States^6^. In Europe alone, approximately 13.5 million people suffer from *Ambrosia*-induced allergies, costing ~7.4 billion euros annually^7^. Continued invasion and climate change are predicted to more than double sensitization to *Ambrosia* pollen^8^, further magnifying the health burden of this pest. Pollen monitoring has demonstrated that climate change has already significantly lengthened the ragweed pollen season, particularly at high latitudes^9^. Consequently, there is considerable incentive to understand the factors that contribute to *Ambrosia* pollen production, including the species’ invasive spread, the timing of pollen production, plant size, and fecundity.

*Ambrosia artemisiifolia* populations are characterized by strong local adaptation and high gene flow between populations^10,11^. In Europe, invasive populations have been established through multiple genetically diverse introductions from North America over the last ~160 years^11–13^. Remarkably, latitudinal clines observed for multiple traits in the native range, including flowering time and size, have re-evolved in Europe and Australia^14^, suggesting rapid local adaptation following invasion. Moreover, this trait-level parallelism is echoed in signals of parallelism at the genetic level^10^.

As biological invasions continue to increase worldwide^15^ and the effects of anthropogenic climate change intensify, understanding the genetic architecture of adaptation to sudden environmental shifts—a classical question in evolutionary research—becomes ever more important. While long-standing theory suggests that evolution in response to incremental environmental change should proceed through mutations of infinitesimally small^16^ or moderate^17^ effect, large-effect mutations are predicted to be useful in bridging extreme, sudden environmental shifts^18^. Moreover, alleles of large effect will, in cases of local adaptation, be better able to persist in the face of the swamping effects of gene flow^19^. Large-effect mutations are also more likely to produce patterns of evolutionary repeatability, or genetic parallelism, between species’ ranges^20^, particularly if adaptive responses make use of standing genetic variation (as would be expected during a bout of rapid adaptation), rather than *de novo* mutations^21^. These features of large-effect mutations may, however, be achieved by groups of mutations in tight genetic linkage^19^, including mutations captured by chromosomal inversions^22^. There is substantial empirical evidence for the involvement of inversions in local adaptation^23^. For example, *Drosophila melanogaster*’s *In(3R)Payne* inversion shows parallel environmental associations across multiple continents^24^, and several plant inversions have been identified as contributing to local adaptation and ecotype divergence^25,26^. Theory also predicts that inversions can drive range expansions^27^, though their actual contributions to biological invasions are not well-understood.

Here we develop a chromosome-level phased diploid reference assembly, and examine genome-wide variation in over 600 modern and historic *A. artemisiifolia* samples from throughout North America and Europe^11^. Using this data of unparalleled spatial and temporal resolution, we first identified regions of the genome experiencing climate-mediated selection in the native North American and introduced European ranges leveraging landscape genomic approaches and genome-wide association studies of adapting traits such as flowering time. Second, motivated by evidence that European and North American populations show similar trait clines with respect to climate^14^, we examined the extent of between-range parallelism at the genetic level. Although adaptive traits such as flowering time and size are polygenic, we expected to see substantial levels of parallelism if large and moderate effect standing variants were contributing to adaptive divergence. Third, we determined if these same regions show temporal signatures of selection in Europe, which would be expected if some invading populations were initially maladapted to their local climates. We coupled this temporal genomic analysis with a temporal analysis of phenological trait changes in European herbarium samples to further support our findings of genomic signatures of selection on flower time genes. Finally, we identified haplotype blocks with multiple features consistent with inversions, in genomic regions enriched for signatures of parallel adaptation. Four of these colocalize with inversions identified in our diploid assembly, confirming their identity as inversions. To determine if these haploblocks were contributing to rapid local adaptation in Europe, we assessed spatial and temporal changes in their frequency as well as their associations with adaptive traits.

## RESULTS

### Reference genome assembly

We assembled a chromosome-level phased diploid *Ambrosia artemisiifolia* reference genome (fig. 1; fig. S1) from a heterozygous, diploid individual collected from Novi Sad, Serbia. After scaffolding with HiRise^28^, our final assembly consisted of two haploid assemblies with genome sizes of 1.11 and 1.07Gbp (flow cytometry estimates of genome size range from 1.13-1.16Gbp^29,30^), with 94.3 and 96.5% of each respective genome assembled into 18 large scaffolds (table S1; table S2), consistent with the 18 chromosomes of *A. artemisiifolia*. Complete copies of all 255 Benchmarking Universal Single-Copy Orthologs (BUSCO^31^) genes were identified on the 18 chromosomes of each haploid assembly, with 183 (71.8%) single-copy and 72 (28.2%) duplicated (fig. 1C; table S1). These high numbers of duplicated BUSCO genes likely reflect the whole-genome duplications experienced in the Asteraceae, including a recent one shared by *Helianthus* (sunflower) species at the base of the tribe^32,33^. This species also retained a large number of duplicated BUSCO genes^33^. A large fraction of the genome consisted of repetitive sequence (67%; table S3). Retroelements were the largest class (39.5%), with long terminal repeats, particularly Gypsy (7.87 %) and Copia (18.98 %), being the most prevalent. MAKER^34^ identified 36,826 gene models with strong protein or transcript support, with average coding lengths of 3kbp and 5.75 exons per gene (fig 1A; table S4).

**Figure 1.**
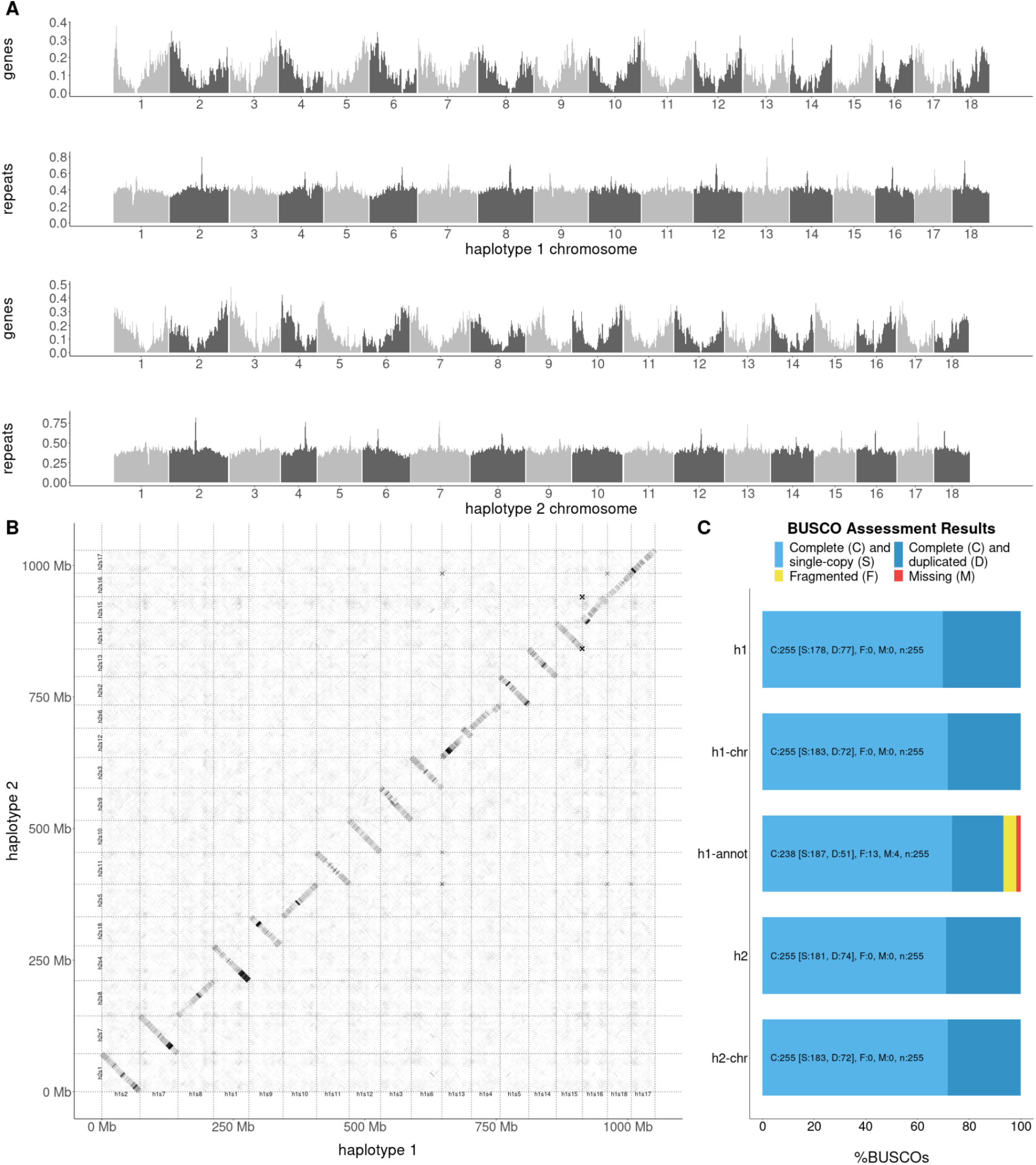
A phased diploid genome assembly of *Ambrosia artemisiifolia*. **A**. The distribution of gene and repeat density in 1Mbp windows across the 18 chromosomes of each haplotype. **B**. Alignments of the 18 chromosomes of each haplotype. **C**. Benchmarking Universal Single-Copy Orthologs (BUSCO) results for each haplotype assembly, each chromosome-only assembly, and the gene annotations for haplotype 1 using the *eukaryota odb10* gene set.

### Genome-wide association studies

Genome-wide association studies (GWAS) using 121 modern samples across North American (*n* = 43) and European (*n* = 78; table S5) ranges identified significant associations with 16 of 30 phenotypes, many of which are putatively adaptive, previously measured by van Boheemen, Atwater and Hodgins^14^ (fig. S2; table S6). All phenotypes yielded associations within at least one predicted gene, including an association between three flowering time phenotypes and a nonsynonymous SNP in the *A. artemisiifolia* homologue of *A. thaliana* flowering-time pathway gene *early flowering 3* (*ELF3*)^35^, an “evolutionary hotspot” for parallel flowering time adaptation in *A. thaliana*, barley and rice^36^ (fig. 2A;B; table S6). Candidate SNPs in *ELF3* are restricted to high-latitude populations in both ranges, where they occur at moderate to high frequencies (fig. 2C). While the latitudes of these populations are greater in Europe than North America, the climatic conditions are similar (fig. S3), indicative of local climate adaptation in parallel between ranges. Additionally, a haplotype containing four nonsynonymous SNPs in an S-locus lectin protein kinase gene was associated with several traits including flowering end date and maximum height (fig. S2; table S6).

**Figure 2.**
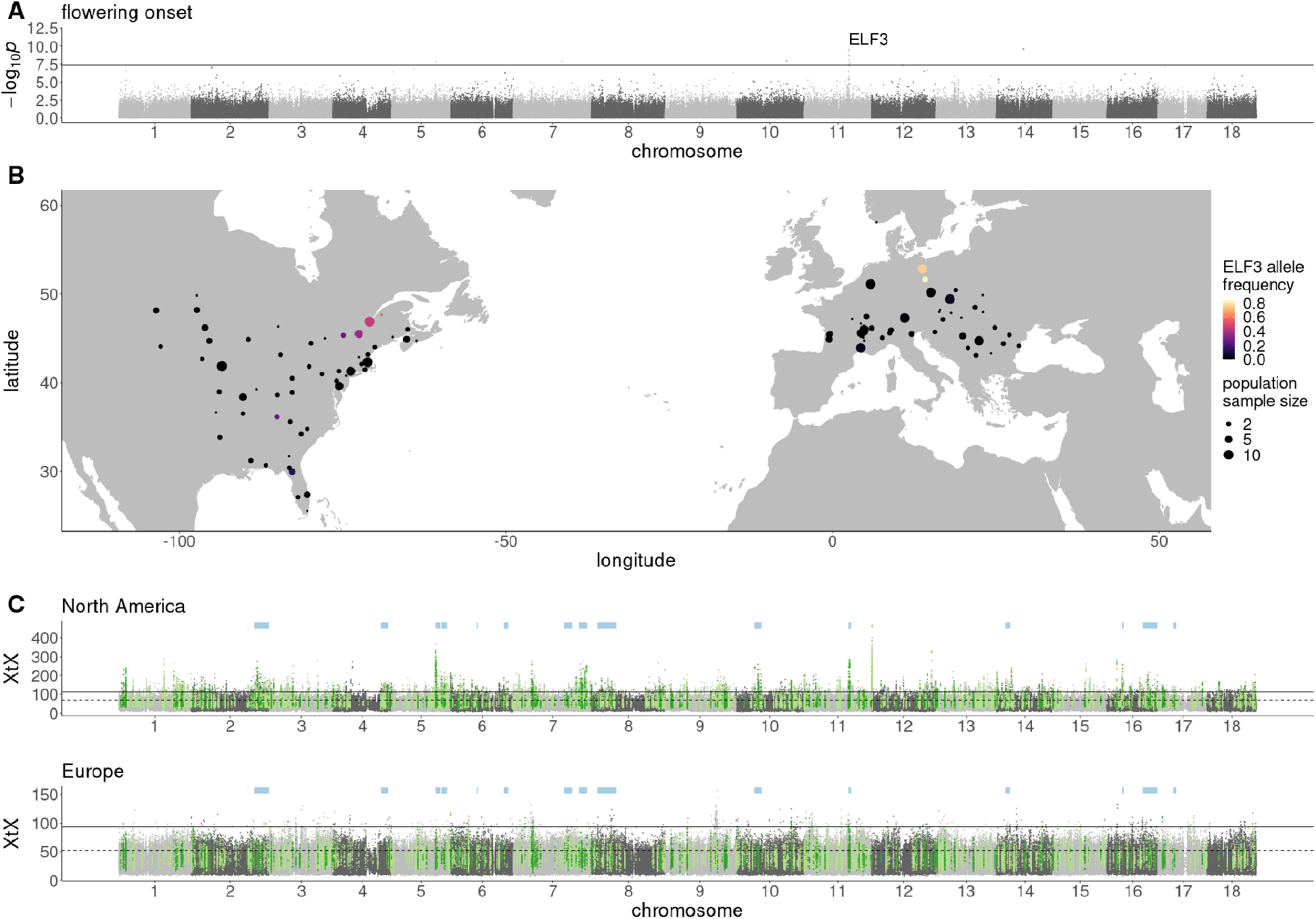
Signatures of climate adaptation in *Ambrosia artemisiifolia*. **A**. GWAS results (−log_10_*p* against genomic location) for flowering onset (solid line indicates a Bonferroni-corrected p-value of 0.05). **B**. Distribution of a strongly-associated nonsynonymous SNP in *ELF3* among modern *A. artemisiifolia* populations used in this study. **C**. Genome-wide XtX scans between sampling locations within each range separately. Solid lines indicate Bonferroni-corrected significance derived from XtX *p*-values; dashed lines indicate the top 1% of genome-wide XtX values. Green highlights represent the top 5% of 10kbp WZA windows for each scan that are also among the top 5% of EAA WZA windows for at least one environmental variable, with dark green indicating outlier windows shared between North America and Europe. Pale blue bars indicate the location of 15 haploblocks (putative chromosomal inversions) that overlap shared outlier windows.

### Environmental-allele associations

To identify genome-wide spatial signatures of local adaptation in North American and European ranges, we performed genome scans for population allele frequencies among *A. artemisiifolia* modern samples that were both highly divergent between populations (BayPass XtX)^37^, and correlated with 19 WorldClim temperature and precipitation variables (table S7)^38^. Statistics were analyzed in 10kbp windows using the Weighted-Z Analysis (WZA)^39^. In North America (143 samples; 43 populations), 2,167 (80.1%) of the 2,704 outlier windows for genomic divergence (XtX) were also outlier windows for at least one environmental variable (XtX-EAA), while in Europe (141 samples; 31 populations), only 1,357 (50.3%) of the 2,697 XtX outlier windows overlapped environmental variable outlier windows. Signatures of local adaptation were much stronger in North America than Europe, with the North American range showing more extreme XtX values (fig. 2C), as well as more XtX-EAA windows (fig. 2C; table S7). This suggests that North American *A. artemisiifolia* exhibits greater population differentiation, and a stronger relationship between population differentiation and the environment than Europe, which is consistent with the expectation that populations from the native range will be better-adapted to their environment than those from the recently-invaded European range.

Previous studies in *A. artemisiifolia* have identified signatures of repeatability between native and invasive ranges at phenotypic and genetic levels^10,14^. We observed congruent patterns in our data: among North American and European XtX-EAA outlier windows, 291 showed parallel associations with the same environmental variable between ranges (significantly more than would be expected by chance; hypergeometric *p* = 1.07×10^−126^; fig. 2C), with 21.4% of climate adaptation candidates in Europe also candidates in North America. To account for the possibility that the number of parallel windows is inflated by extended linkage disequilibrium between windows (and hence represents a smaller number of loci), we combined consecutive outlier windows, and windows in haploblock regions, into single windows and repeated the analysis, in which the parallelism remained highly significant (hypergeometric *p* = 1.29×10^−91^). Consequently, many of the same regions of the genome are involved in climate adaptation in both ranges.

North American, European and parallel XtX-EAA outlier windows included 28, 22, and three flowering-time pathway genes, respectively, however this only represented a significant enrichment (Fisher’s exact test *p* < 0.05) in North America (table S8). GWAS flowering time candidate *ELF3* was located in a parallel XtX-EAA window, while the flowering and height-associated S-locus lectin protein kinase gene was in a North American XtX-EAA window only. Gene ontology terms enriched in parallel XtX-EAA windows included “iron ion binding” and “heme binding” (terms relating to cytochrome P450 genes), as well as “gibberellin biosynthetic process” (table S9). Some cytochrome P450 genes are involved in detoxification of xenobiotic compounds and the synthesis of defense compounds, while others play key developmental roles^41^, including contributing to the biosynthesis of gibberellin, a hormone that regulates a range of developmental events including flowering^42^.

### Temporal phenotypic analysis

To further identify the features of recent adaptation in *A. artemisiifolia*, we leveraged herbarium samples, collected from as early as 1830, for phenotypic and genomic analyses. A trait-based analysis of 985 digitized herbarium images (fig. S4) identified a significant shift in the probability of flowering and fruiting over time in Europe, but this change depended on the latitude or the day of the year the sample was collected (fig. S5; table S10). For the trait presence of a mature male inflorescence, we identified a significant interaction between collection year and latitude (*F_1,886_* = 7.89, *p* < 0.01) where in northern populations, more recently collected plants were more likely to be flowering than older historic specimens. For this trait, collection day also significantly interacted with collection year and more recently collected plants were more likely to be flowering later in the year and less likely to be flowering earlier in the year. Similar patterns were identified with the presence of fruit, as older samples were less likely to produce fruit later in the season compared to recent samples (day-by-year interaction *F*_1,886_ = 32.33, *p* < 0.001; fig. S5; table S11). This substantial spatio-temporal change in phenology is consistent with experimental common gardens that show that earlier flowering has evolved in northern populations and later flowering in southern populations following the invasion of Europe^14^. Further, this shift in both flowering and fruit set over time supports the hypothesis that an initial mismatch between the local environment and the genotypes present impacted the reproductive output of *A. artemisiifolia* during the early stages of colonization, particularly in northern Europe.

### Genomic signatures of local selective sweeps

Genome resequencing of *A. artemisiifolia* herbarium samples^11^ allowed comparisons between historic and modern populations. We grouped historic samples based on their age and proximity to a modern population sample, resulting in five North American and seven European historic-modern population pairs (table S12), which were scanned for signatures of local selective sweeps by identifying windows with extreme shifts in allele frequency and extreme reductions in diversity over time. We found far more evidence for recent sweeps in Europe (476 unique windows) than in North America (129 unique windows; fig. S6 c.f. S7; table S12), consistent with the expectation that a haphazardly-introduced invader will frequently be maladapted, initially, and undergo rapid adaptation to local environmental conditions following its introduction. The most dramatic selective sweep signature was observed in Berlin over a ~14Mbp region on chromosome 2 (fig. 3A; fig. S7), which accounts for 274 (58%) of the European sweep windows. In Berlin, sweep windows were also observed containing and surrounding the flowering onset GWAS peak that includes *ELF3*. Scans of Fay and Wu’s *H* in modern populations provide further evidence for recent selection in these regions, and suggest more geographically widespread selection in the region on chromosome 2 (fig. S8; S9). In comparisons of spatial and temporal signatures of selection, three and five sweep windows were also XtX-EAA outliers in North America and Europe respectively. All such windows in Europe were in the chromosome 2 or *ELF3* regions. To further investigate the temporal shift associated with *ELF3*, we focussed on the nonsynonymous *ELF3* variant across all samples within a 200km radius of Berlin (15 historic and eleven modern samples). The frequency of the variant increased from 2.6% to 73.9% between historic and modern samples, an allele frequency shift greater than 10,000 putatively neutral loci sampled from the same geographic region; similar shifts were not observed in North American samples within a 200km radius of Quebec City (ten historic and eleven modern), where the *ELF3* allele is at comparably high frequencies (fig. 3C).

**Figure 3.**
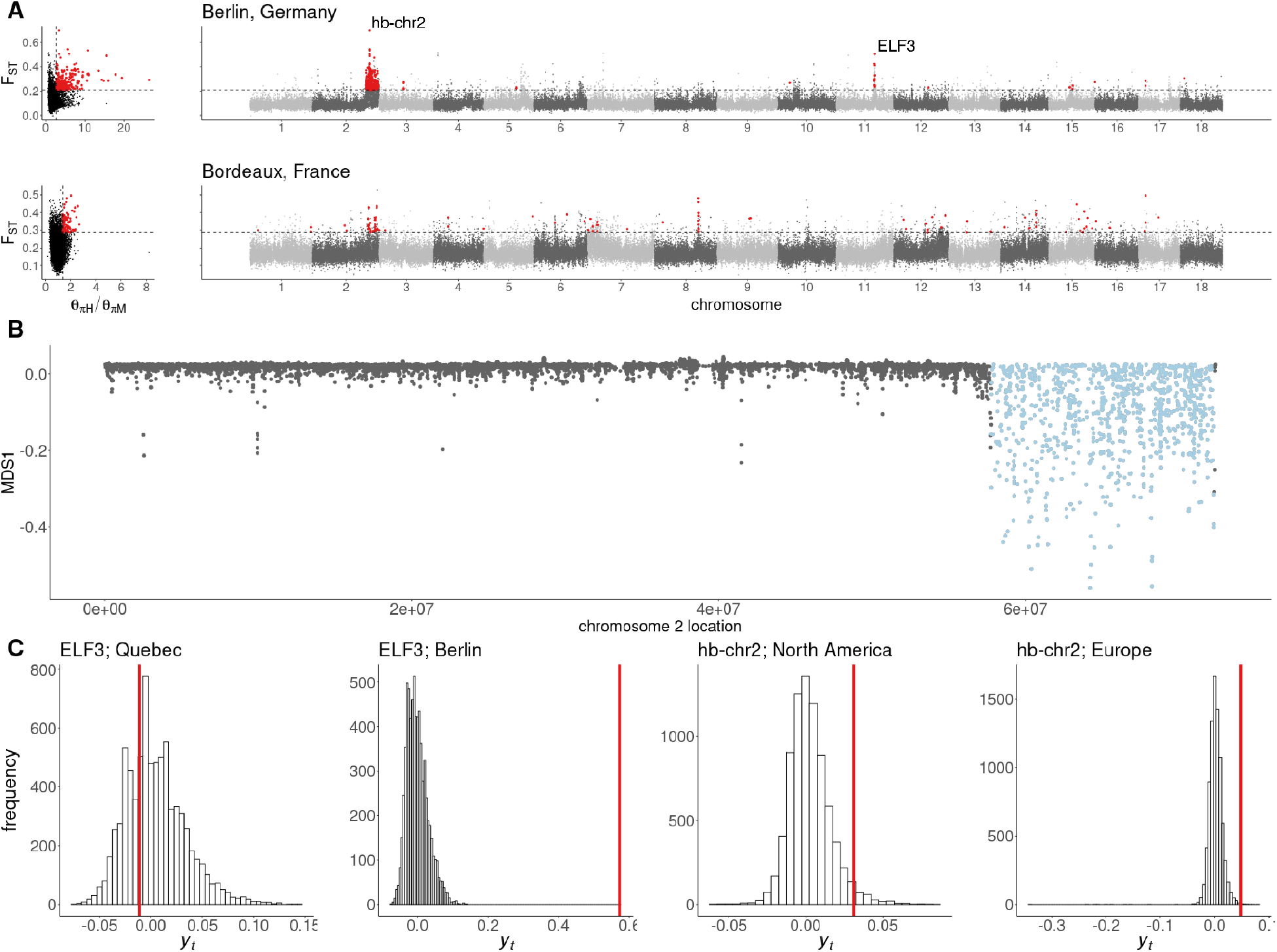
Temporal signatures of selective sweeps in Europe. **A**. Distributions of F_ST_ between historic and modern samples and the ratio of historic to modern nucleotide diversity (θ_πH_/θ_πM_) from Berlin and Bordeaux, and F_ST_ against genomic location. Red points indicate putative selective sweep windows, which are in the top one percent of per-window F_ST_ and θ_πH_/θ_πM_ (dashed lines). **B**. Strong evidence for a selective sweep on chromosome 2 in European populations corresponds with local divergent population structure (MDS1), indicating the presence of a haploblock (putative chromosomal inversion) in this region. **C**. A standardized measure of allele frequency change, *y_t_* (calculated according to equation 1) for shifts between historic and modern populations across putatively neutral SNPs (histograms) and selective sweep candidates (red lines).

### Haploblock identification

Chromosomal inversions have previously been identified as driving local adaptation of ecotypes of *Helianthus* species^26^. We used a similar approach to identify genomic signatures of putative inversions (haploblocks) contributing to local adaptation in *A. artemisiifolia*. Briefly, we identified genomic regions in which population structure was divergent and fell into three clusters, putatively representing the heterozygous and two homozygous genotypic classes of an inversion. Further, we looked for pronounced shifts in population structure (indicating inversion breakpoints), elevated local heterozygosity in the heterozygous cluster, and increased linkage disequilibrium across the region (fig. S10). We examined mapping populations of *A. artemisiifolia*^40^ for evidence of map-specific reductions in recombination across haploblock regions (i.e., suppressed recombination in haploblock regions in some maps but not others; fig. S11). This would be the pattern expected when recombination is suppressed by inversions in heterozygotes but not homozygotes, as opposed to the haploblocks being caused by global reductions in recombination in those regions. Most haploblocks with sufficient markers in the region showed evidence of suppressed recombination in some maps but not others, with the exception of hb-chr6b, which showed suppressed recombination in all maps. To validate our haploblock detection, we examined an alignment of our two haploid reference genomes and identified four segregating inversion polymorphisms corresponding to haploblocks (fig 1B; fig. S12).

Focussing our analysis on regions showing signatures of adaptation, we identified 15 haploblocks with the above genomic signatures of chromosomal inversions overlapping the 291 WZA windows that were parallel outliers for both XtX and at least one climate variable: hb-chr2 (14.5Mbp), hb-chr4 (6.2Mbp), hb-chr5a (4.1Mbp), hb-chr5b (5.4Mbp), hb-chr6a (1.1Mbp), hb-chr6b (3.9Mbp), hb-chr7a (7.3Mbp), hb-chr7b (7.0Mbp), hb-chr8 (17.3Mbp), hb-chr10 (6.5Mbp), hb-chr11 (2.2Mbp), hb-chr14 (4.9Mbp), hb-chr16a (1.7Mbp), hb-chr16b (13.7Mbp) and hb-chr17 (2.8Mbp). These haploblocks contained 77 of the 291 parallel XtX-EAA windows (26.5%; fig. 2C), although they only represent ~10% of the genome (a significant enrichment; hypergeometric *p* = 5.8×10^−17^). One haploblock also corresponds to the European selective sweep region on chromosome 2 (fig. 3A;B). This suggests that these haploblock regions have played a pivotal role in generating parallel signatures of selection observed in *A. artemisiifolia*.

### Haploblock frequency changes through space and time

To identify changes in haploblock frequency over time and space, which would be consistent with selection on these putative inversions, we first estimated haploblock genotypes for all historic and modern samples. Within haploblock boundaries identified using modern sample SNP data in Lostruct^43^, we performed local PCAs with both historic and modern samples (table S4) and identified genotypes by kmeans clustering (fig. S9). For modern samples, we used generalized linear models to estimate the slopes of the haploblock frequencies as a function of latitude within each range. For those haploblocks that were significantly associated with latitude, we compared these estimates with the genome-wide distribution of slopes for North America and Europe, based on 10,000 unlinked SNPs that were randomly selected from outside haploblocks and genes. The estimated slopes for eight haploblocks fell into the 5% tail of the distribution for at least one of the ranges (fig. S13A). However, this approach did not examine temporal changes nor the combined signatures of selection over space and time. To do so, we ran generalized linear models comparing haplotype frequency with latitude, time (date of specimen in years) and range (North America vs. Europe; fig. 4A; fig. S14; table S13-S18). All but two (hb-chr6a and hb-chr7a) of the haploblocks showed significant changes over time, space or both time and space. These patterns were robust to time being coded as discrete (historic vs. modern) or as continuous (by year; table S13). Most showed temporal changes either in their average frequency in one or both ranges, or in their relationship with latitude within each range, a pattern that is consistent with recent local selection on these haploblocks. Most of these haploblocks also showed significant associations with latitude in at least one range or timepoint, indicative of climate adaptation. For instance, hb-chr10, hb-chr14, hb-chr16b and hb-chr17 all showed significant parallel latitudinal associations in both ranges in historic and modern samples. For hb-chr5b, haplotype frequency was negatively correlated with latitude in modern and historic samples from North America, as well as in modern European samples, consistent with climate-mediated selection. However, historic European populations did not display an association with latitude, which may reflect maladaptation during the initial stages of the European range expansion (fig. 4A; table S16, S18). In one case (hb-chr7b) the significant latitudinal clines showed opposing yet significant slopes for each range in the modern samples, perhaps indicating contrasting patterns of local selection between the ranges. For hb-chr2 the latitudinal clines formed over time, with a substantial increase in frequency over time particularly at higher latitudes.

**Figure 4.**
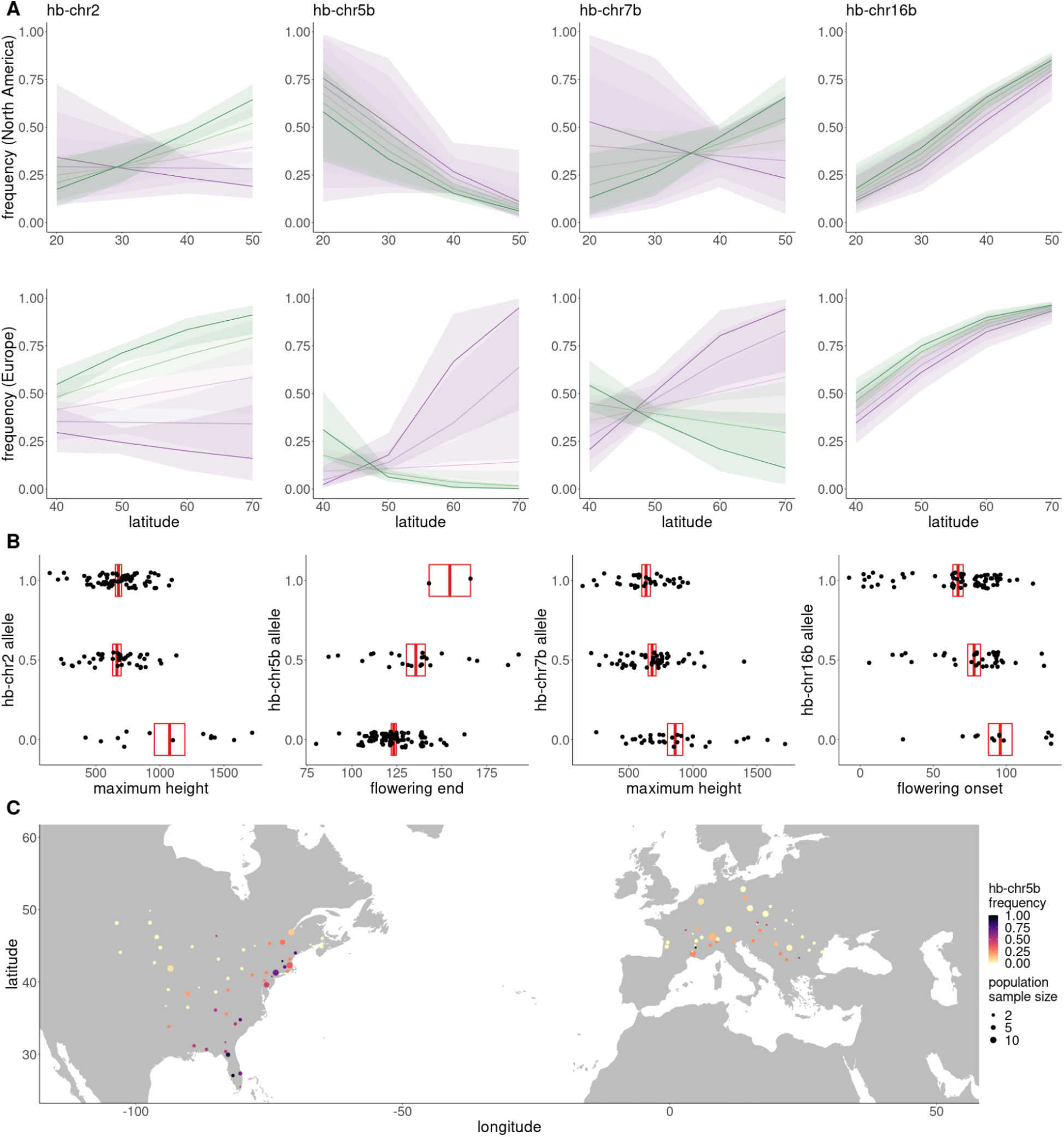
Haploblock distributions and trait associations. **A**. Logistic regression models with 95% CI ribbons (see table S13-18 for model details) of haploblock frequency (allele 1) against latitude for four haploblocks across five time bins ranging from most historic (purple) to most modern (green). **B**. Examples of significant associations between haploblock alleles and phenotypes (boxes denote mean and SEM). **C**. hb-chr5b allele frequency in modern *A. artemisiifolia* populations.

To further investigate whether European populations showed evidence of recent local selection on the haploblocks, we tested whether estimates of selection inferred from contemporary spatial data were associated with temporal changes in haploblock frequencies between historical and contemporary European populations. We used spatial variation in contemporary haploblock frequencies to estimate the relative strength of local selection on these haploblocks (see supplementary text S2). We specifically compared estimates of the maximum slope of latitudinal clines for each putative inversion’s frequency to simple population-genetic models for clines at equilibrium between local selection and gene flow. In these models, cline slopes are proportional to 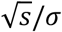, where *s* represents the strength of local selection for a given inversion and *σ* is the average dispersal distance of individuals in the range^44^ (supplementary text S2; table S16). While our estimates of selection are, therefore, scaled by the dispersal rate, dispersal should equally affect all inversions within a given range, allowing us to infer the relative strength of spatially varying selection for each putative inversion. We found that estimates of the relative strengths of local selection across the haploblocks in modern samples were correlated between the ranges, indicating parallel patterns of local selection along the latitudinal gradient (*r* = 0.55, *p* = 0.03). We also found that changes in the haploblock cline slopes between historic and modern time points within Europe were significantly correlated with our estimates of the relative strength of spatially varying selection of the haploblocks across the European range (*r* = 0.86, *p* < 0.001 fig. S13B). Such a pattern is consistent with a scenario in which historical European populations were not initially locally adapted (haploblock frequencies were initially far from local optima) and where the haploblocks subject to relatively strong local selection exhibited the greatest temporal changes in local frequency over the ensuing century. The same pattern was not observed in the North American native range, whose historic populations are likely to have been consistently closer to the local optima across the timescale of our analysis (*r* = 0.25, *p* = 0.36). Cline slopes for latitude were also shallower in historic relative to modern European populations (*t*_10.38_ = −3.09, *p* = 0.01; mean absolute slope: historic EU = 0.05; modern EU = 0.14), but not so in North America (*t*_17.81_ = −0.37, *p* = 0.71), consistent with initial maladaptation in Europe, followed by adaptation to local climates.

The hb-chr2 haplotype frequency dramatically increased in many populations over time, particularly at mid to high latitudes, which encompassed the European populations and northern North American populations (fig 4; tables S13;S18). Signatures of selective sweeps were observed in the hb-chr2 region in European populations in historic to modern comparisons of divergence and diversity (fig. 3A; fig. S7). In this same region, negative outlier values of Fay & Wu’s *H* relative to the genome-wide pattern were identified in three of the five North American and six of the seven European locations, indicating an excess of high frequency derived SNPs and implicating recent positive selection (fig. S8; S9). To further distinguish the effects of drift from those of selection, we compared an empirical null distribution of allele frequency changes over time in Europe using 8,303 SNPs matched for allele frequency (fig. 3C). Only 11 (0.13%) of the 8,303 loci in this null distribution showed an allele frequency change greater than hb-chr2. We then estimated strengths of selection for the putative inversion that are consistent with the observed increase in frequency within the invasive range (see supplementary text S1). With positive selection for the putative inversion, the estimated frequency shift at the center of the European range is consistent with a 2.4% difference in fitness (95% CI = {1.5%, 3.3%}) between individuals homozygous for the inversion relative to homozygotes for the alternative haplotype. Scenarios of balancing selection require stronger selection to explain the inversion frequency shift over time (see supplementary text S1). This estimate is smaller than many empirical estimates of selection on individual loci in natural populations^45^. The greater timespan of our samples facilitated detection of strong signals of selection for loci that would otherwise be missed by the short time periods afforded by most temporal studies.

### Biological functions of haploblocks

Several analyses provide evidence for the biological function of these putatively adaptive haploblocks. From genome annotations, we found that haploblocks were collectively enriched for flowering-time pathway genes (Fisher’s exact test *p* = 0.002), although individually only hb-chr5b was significantly enriched (Fisher’s exact test *p* = 0.001;table S8), consistent with a large-effect flowering time QTL identified within this haploblock^40^. hb-chr2 was enriched for the “recognition of pollen” gene ontology term, with 36 (17%) of the genome’s 210 genes annotated with this term falling within this haploblock (table S9). hb-chr5a was enriched for genes with the “pectate lyase activity” term, including the top BLAST hit for *Amba1* (99.7% identity, *E*-value = 0), which encodes the *A. artemisiifolia* protein responsible for the majority of allergic reactions^46^. A detailed analysis of the hb-chr5a region reveals a cluster of six closely-related pectate lyase genes, which correspond with elevated XtX and XtX-EAA outlier windows in both ranges (fig. S15). hb-chr11 also overlaps the flowering time GWAS candidate *ELF3* (fig. 2A;B), and the nonsynonymous variant which displays strong patterns in GWAS is only observed on one of the haploblock genotype backgrounds. We also identified phenotypic associations with haploblocks by encoding haploblock genotypes into our GWAS pipeline. Significant associations (*p* < 0.05 Bonferroni-corrected for multiple testing across 15 haploblocks) were observed for four haploblocks, including for traits related to flowering time (fig. 4B; table S19).

## DISCUSSION

We have described, at unprecedented temporal and spatial resolution, the evolutionary-genetic changes accompanying a recent and rapid invasion by a noxious pest. Our study system, while unique in many ways, yields results with important general implications for our understanding of the genetic basis of rapid adaptation to environmental change and the pervasiveness of parallel evolution in geographically widespread species.

While invasive species are often envisaged to encounter novel selection pressures as they spread across alien landscapes, they must also readapt to similar environmental variation encountered in their native range, as haphazardly-introduced invaders are unlikely to be well-adapted to local conditions when the invasive populations initially expand across climatic gradients. Much of *Ambrosia artemisiifolia*’s European invasion lies within climatic extremes encountered across its native range (fig. S3). Despite this similarity in climatic variation, the patterns of parallel climate adaptation between native and invasive ranges are striking given the evolutionarily recent introduction of the species into Europe. As *A. artemisiifolia*’s invasion of Europe consisted of multiple introductions over a brief evolutionary time scale, these patterns are likely examples of ‘collateral evolution’^47^, in which standing genetic variation in *A. artemisiifolia*’s native range has been co-opted for adaptation in and across the European invasive range. Parallel evolution is a hallmark of natural selection and parallel changes at the genetic level point to constraints and biases in the genetic pathways to adaptation that are evolutionarily achievable; when certain paths to adaptation are favored, such as when beneficial variants are already present in the population as standing variants, evolution will repeatedly draw on the same subset of genes to reach the same adaptive endpoints.

From herbarium specimens that were sampled throughout the course of *A. artemisiifolia*’s invasion of Europe, we observed an abrupt change in flowering and fruiting over time. Leveraging whole-genome sequences of herbarium samples across North America and Europe, we were also able to scan populations for temporal genomic signatures of selective sweeps. Although some populations have experienced shifts in ancestry over time in Europe^11^, peaks against the genome-wide background provide compelling evidence for rapid local adaptation in European populations, with the strongest genetic signals of rapid change over time corresponding to some of the strongest signatures of local adaptation in our spatial analyses, particularly windows in the region of the *ELF3* gene and hb-chr2. Further, these regions show parallel signals of climate adaptation in North America and are associated with adapting traits such as flowering onset. These multiple lines of evidence provide strong support that climate-mediated selection on phenology was pivotal in shaping the adaptive genetic landscape of *A. artemisiifolia* in Europe.

Large haploblocks (putative inversions) contribute substantially to these genetic signals of parallel adaptation. 27% of these haploblocks correspond to inversions segregating in our diploid assembly. We propose that these haploblocks maintain cassettes of co-selected genes that effectively segregate as single alleles of large effect^22,27^, providing a genetic architecture suited to local adaptation in the face of high gene flow^11,19^. Consistent with this hypothesis, haploblocks are enriched for genes with particular biological functions, display associations with locally-adaptive traits, and carry signals of strong selection in both the native and invasive ranges. The evolution of inversions along environmental gradients has been reported in a range of species^23^. However, by investigating haploblocks in an invasive plant with extensive timestamped collections, we have demonstrated dramatic and adaptive evolutionary change of inversions under natural conditions, providing compelling evidence of strong and recent natural selection. These data have also allowed us to estimate selection for these variants, and we have shown that haploblocks with the strongest estimates of clinal selection are driven more rapidly towards their putative equilibria within the invasive range.

An important question during this era of environmental upheaval is the role of adaptation during range expansion and its necessity during colonization. Through our analysis of historic samples, we have shown that *A. artemisiifolia* was present in regions throughout Europe well before many of these adaptive variants became locally common, suggesting the species’ extensive phenotypic plasticity may have facilitated its initial expansion. Strong local selection further improved the match between genotypes and local environments, even appearing to affect reproductive output in herbarium specimens. Many of the selected variants we identified are linked to traits that are key factors in the timing, length and severity of the local pollen season (e.g. days to flowering onset, days to the end of pollen production, and biomass). Consequently, local adaptation has played a central role in shaping the allergy season in Europe and will likely continue to be critical as climate change and continued range expansion further amplify the damaging effects of this hazardous weed^48^.

## METHODS

### Genome assembly

Seeds collected from a wild *Ambrosia artemisiifolia* population in Novi Sad, Serbia (lat. 45.25472, lon. 19.91231) were sown in potting soil at a greenhouse facility at the Ringve Botanical Garden, NTNU University Museum (Trondheim. Norway). After 160 days of growth under stable light and watering conditions, young leaf tissue from mature individual plant “NSS02/B” was sampled and flash-frozen in liquid nitrogen. These tissues were then shipped to Dovetail Genomics for high molecular weight DNA extraction and library building.

DNA samples were quantified using Qubit 2.0 Fluorometer (Life Technologies, Carlsbad, CA, USA). The PacBio SMRTbell library (~20kbp mean insert length) for PacBio Sequel was constructed using SMRTbell Express Template Prep Kit 2.0 (PacBio, Menlo Park, CA, USA) using the manufacturer recommended protocol. The library was bound to polymerase using the Sequel II Binding Kit 2.0 (PacBio) and loaded onto PacBio Sequel II). Sequencing was performed on PacBio Sequel II 8M SMRT cells generating 65.9Gbp of data. These PacBio CCS reads were used as an input to Hifiasm^49^.

For each Dovetail Omni-C library, chromatin was fixed in place with formaldehyde in the nucleus and then extracted. Fixed chromatin was digested with DNAse I, chromatin ends were repaired and ligated to a biotinylated bridge adapter followed by proximity ligation of adapter containing ends. After proximity ligation, crosslinks were reversed and the DNA purified. Purified DNA was treated to remove biotin that was not internal to ligated fragments. Sequencing libraries were generated using NEBNext Ultra enzymes and Illumina-compatible adapters. Biotin-containing fragments were isolated using streptavidin beads before PCR enrichment of each library. The library was sequenced on an Illumina HiSeqX platform to produce ~30x sequence coverage. The PacBio CCS reads and Omni-C reads were then used as input for Hifiasm to produce two haplotype-resolved assemblies (hap1 and hap2) using default parameters.

HiRise was used (see read-pair above) to scaffold each haplotype-resolved assembly. Each *de novo* assembly and Dovetail OmniC library reads were used as input data for HiRise, a software pipeline designed specifically for using proximity ligation data to scaffold genome assemblies^28^. Dovetail OmniC library sequences were aligned to the draft input assembly using bwa^50^. The separations of Dovetail OmniC read pairs mapped within draft scaffolds were analyzed by HiRise to produce a likelihood model for genomic distance between read pairs, and the model was used to identify and break putative misjoins, to score prospective joins, and make joins above a default threshold (fig. S1C). The NCBI^51^ genome submission portal identified 30 (7.4Mbp total) and 26 (6.4Mbp total) scaffolds in haplotypes 1 and 2 respectively containing bacterial contamination which were subsequently removed from the final assembly.

We used GenomeScope 2.0 to estimate the genome size and ploidy using 21mers identified in the reads with Jellyfish 2.3.0^52^. Genomescope estimated the haploid genome size to be 1.04Gbp using a diploid model (fig. S1A), a better model fit (95%) than the tetraploid model (91%), which also vastly underestimated the haploid genome size (497 Mbp). This finding was consistent with the smudgeplot produced by Genomescope, which also indicated diploidy (fig. S1B). The final assembly sizes for each haplotype were 1.11 and 1.07Gbp, which were similar to the GenomeScope estimates. BUSCO (Benchmarking Universal Single-Copy Orthologs) version 5.1.3^31^ analysis of each assembly using the eukaryota odb10 dataset (table S1) demonstrated that both assemblies were complete with relatively low levels of duplication given the history of whole genome duplication in the tribe.

To assess the presence of remnant haplotigs and other assembly artifacts, we mapped Illumina reads used in the reference genome assembly to haplotype 1 of the reference genome using AdapterRemoval^53^, BWA-MEM^50^ and Picard MarkDuplicates (https://broadinstitute.github.io/picard/), and measured average sequencing depth and heterozygosity of the alignment in non-overlapping 1Mbp windows across the genome. Window depth was never greater than two times higher or 0.5 times lower than the mean, and furthermore regions of both low depth and low heterozygosity were distributed throughout the genome. The fact that there were no large regions with both low read-depth and low heterozygosity points to the success of the haplotype-resolved assembly (fig. S1D). Minimap2 was used to align each haplotype against itself and against one another, after filtering for alignments shorter than 10kbp and with fewer than 5000 matches, to identify homologous blocks that may represent haplotigs. This analysis revealed the presence of a misassembly where each assembly contained a region of scaffold 18 duplicated on scaffold 19, while the orthologous region was missing in the alternative assembly. This suggested that a section of scaffold 18 in each haplotype-resolved assembly had been incorrectly placed in the wrong haplotype (corresponding to chromosome 18 in haplotype 1 and chromosome 9 in haplotype 2). After making these manual corrections, the genetic map confirmed the continuity of these chromosomes in each haplotype (fig. S16). The alignments of the final corrected assemblies within and between the assemblies further confirmed the continuity of the assemblies and the absence of haplotigs (fig. 1B).

### Whole-genome resequencing samples

Whole-genome resequencing data used in this study have previously been described in Bieker *et al.^11^*. Modern samples were field-collected between 2007 and 2019, and historic samples were sequenced from herbarium specimens collected between 1830 and 1973. 121 modern samples with corresponding phenotype data collected by van Boheemen, Atwater and Hodgins^14^ were used for genome-wide association studies. 284 modern samples (from populations with a sample size >= 2) were used for environmental-allele associations. 97 historic and 100 modern samples divided into twelve populations were used for historic-modern population comparisons (table S12). For *ELF3* analysis, 26 samples from within 200km of Berlin (15 historic and eleven modern) and 21 samples within 200km of Quebec City (ten historic and eleven modern) were used. Genotyping and analysis of haploblocks was performed using 311 modern and 305 historic samples. For details of each sample see table S5.

### Sample alignment, variant calling and filtering

FASTQ files from historic and modern *A. artemisiifolia* samples from North America and Europe^11^ were aligned to haplotype 1 of our new reference genome using the Paleomix pipeline^54^, which incorporates AdapterRemoval^53^, BWA-MEM^50^, Picard MarkDuplicates (https://broadinstitute.github.io/picard/) and GATK IndelRealigner^55^. Mean depths of alignments ranged from 0.37X to 19.95X with a mean of 4.05X for historic samples, and 1.75X to 44.03X with a mean of 6.86X for modern samples (table S5). Variants were called in the higher-depth modern samples using GATK UnifiedGenotyper^56^ on all contigs greater than 100kbp in length. GATK VariantFiltration^55^ and VcfTools^57^ were used to filter variant calls. SNP and indel calls were separately filtered using GATK hard-filtering recommendations (SNPs: QD < 2.0, FS > 60.0, SOR > 3.0, MQ < 40.0, ReadPosRankSum < −8.0, MQRankSum < −12.5; indels: QD < 2.0, FS > 200.0, SOR > 10.0, ReadPosRankSum < −20.0, InbreedingCoeff < −0.8). Additionally, SNPs and indels were separately filtered for sites with depth (DP) less than one standard deviation below the mean, and greater than 1.5 standard deviations above the mean. Individual genotypes were set to missing if their depth was less than three, then variants with greater than 20% missing across all samples were removed. Samples with greater than 50% missing variants were removed. For the remaining 311 modern samples, genotypes were phased and imputed using Beagle 5.2^58^.

### Genome annotation

To obtain RNA transcript sequences for annotation of the genome, after 160 days of growth additional samples of leaf, stem, flower, root, and branch were taken from individual “NSS02/B” and flash-frozen in liquid nitrogen. From these we extracted RNA from seven tissues (young leaf, old leaf, stem, branch, and three stages of development of the floral head) using a Spectrum Plant Total RNA Kit (Sigma, USA) with on-column DNA digestion following the manufacturer’s protocol. RNA extracts from all five tissues were pooled into a single sample. mRNA was enriched using oligo (dT) beads, and the first strand cDNA was synthesized using the Clontech SMARTer PCR cDNA Synthesis Kit, followed by first-strand synthesis with SMARTScribeTM Reverse Transcriptase. After cDNA amplification, a portion of the product was used directly as a non-size selected SMRTbell library. In parallel, the rest of amplification was first selected using either BluePippin or SageELF, and then used to construct a size-selected SMRTbell library after size fractionation. DNA damage and ends were then repaired, followed by hairpin adaptor ligation. Finally, sequencing primers and polymerase were annealed to SMRTbell templates, and IsoSeq isoform sequencing was performed by Novogene Europe (Cambridge, UK) using a PacBio Sequel II instrument, yielding 97,819,215 HiFi reads. To prepare the raw IsoSeq RNA data for downstream use in the annotation of the genome, we first identified the transcripts in the PacBio single-molecule sequencing data by following the IsoSeq v3 pipeline provided by PacificBiosciences (https://github.com/PacificBiosciences/IsoSeq). Briefly, the pipeline takes PacBio subread files as an input and undergoes steps of consensus generation, demultiplexing of primers, IsoSeq3 refinement, followed by a final clustering of the reads.

Prior to annotation of the genome, repetitive elements were identified using RepeatModeler2^59^. ProExcluder^60^ was then run to remove any protein coding genes from the repeat library. RepeatMasker^61^ was used to mask the genome using the finalized repeat library (table S3). A large fraction of the genome consisted of repetitive sequence (66.51%; fig. 1A). Retroelements were the largest class (39.13%), with long terminal repeats, particularly Gypsy (7.73 %) and Copia (18.82 %), the most prevalent retroelements.

Genome annotation was performed using the MAKER v.3.01.03 ^62^ pipeline. Genome assembly fasta file, the custom repeat library, IsoSeq clustered reads merged with a previously described transcriptome^11^ (as expressed sequence tag [EST] evidence) and protein homology evidence from a plant protein database which combines the Swissprot plant protein database and NCBI Refseq for plants excluding transposable elements were used as the input files for the first run of the annotation pipeline. The custom repeat library was used to mask the repetitive regions. Additional regions with low complexity were soft masked using RepeatMasker v.4.1.1 ^61^.Gene predictors SNAP v.2013-11-29 ^63^ and AUGUSTUS v.3.3.3 ^64^ were trained by running iterative runs of Maker as recommended by ^62^.As the first round of annotation was based on the alignment of the EST evidence to the genome, est2genome option in the Maker control file was set to one to allow Maker to infer gene models directly from the EST evidence. After the completion of the first round of annotations, gene models with an AED (Annotation Edit Distance) score of 0.25 or greater and a length of 50 or more amino acids were retained and used to train SNAP v.2013-11-29 ^63^ to obtain a SNAP hmm file. We then trained AUGUSTUS v.3.3.3 ^64^ using BUSCO v.3.0.2 ^65^. First, training sequences were identified using the gene models predicted by Maker from the first run by excising regions with mRNA annotations and 1000 bp on either side. These were used to run BUSCO using the embryophyte set of conserved genes. After training both SNAP and Augustus, Maker was run again, with SNAP hmm and Augustus files. A total of three rounds of training for each gene predictor were run. We used the script genestats ^66^ to calculate the numbers and lengths of genes, exons, introns and UTR (untranslated region) sequences present in the predicted gene models by the final Maker run. We ran BUSCO v.5.1.3 ^31^ with the eukaryota_odb10 lineage data set on the predicted transcript fasta file by Maker to assess the quality and the completeness of the annotated genome.

For haplotype 1, a high confidence gene set of 36,826 gene models with strong protein or transcript support was identified (table S4). Gene models were compared with *Arabidopsis thaliana* annotations (TAIR10 representative gene model proteins^67^) and the UniProtKB plants database using the *blastp* command in BLAST+^68^. Using an *E*-value threshold of 1×10^−6^, 32,370 (87.9%) genes matched TAIR10 annotations and 28,092 (76.3%) matched UniProtKB. We identified 98.4% of the core eukaryotic genes amongst our annotated genes, 73.3% being single copy, 20% being duplicated and 5.1% fragmented compared to BUSCO markers present in the library *eukaryota_odb10.2020-09.10* (fig. 1C). Gene ontology (GO) enrichment was assessed using GO terms from *A. thaliana* TAIR 10^67^ BLAST results. To identify GO terms enriched among candidate lists, the R/topGO package^69^ was used with Fisher’s exact test, the ‘weight01’ algorithm, and a *p*-value < 0.05 to assess significance. Additionally, annotations were cross-referenced with 306 *A. thaliana* FLOR-ID flowering time pathway genes^70^. 513 predicted *A. artemisiifolia* genes were matched to this dataset, representing 218 unique FLOR-ID genes. Enrichment of flowering time genes was also assessed in candidate gene lists using Fisher’s exact test and a *p* < 0.05 threshold. The effects of imputed variants on predicted genes were estimated using SnpEff^71^.

### Allele frequency outliers and environmental allele associations

Imputed genotype data from modern samples were divided for between-range and within-range analyses in PLINK 1.9^72^, and a minor allele frequency threshold of 0.05 was applied within data subsets. For within-range analyses, sampling locations with fewer than two samples were excluded and allele frequencies were calculated for each sampling location, resulting in 1,150,328 SNPs across 143 samples and 43 populations in North America and 1,132,342 SNPs across 141 samples and 31 populations for Europe. Allele frequency outliers were identified within each range using the BayPass core model^37^, with an Ω covariance matrix computed from 10,000 randomly-sampled SNPs that were located outside annotated genes and haploblocks, and pruned for linkage disequilibrium using a window size of 50kb, a step size of 5bp and an *r^2^* of 0.5 in PLINK^72^. To identify allele frequency variation associated with environmental variables within ranges, 19 bioclimatic variables were extracted for each sampling location from the WorldClim database^38^ using the R/raster package^73^. Population allele frequencies were assessed for correlation with 19 bioclimatic variables using Kendall’s τ statistic in R^74^. Genome-wide XtX and τ results were analyzed in non-overlapping 10kbp windows using the weighted-Z analysis (WZA)^39^, with the top 5% of windows designated outliers.

### Genome-wide association studies

Imputed genotypes from modern samples were filtered in PLINK 1.9^72^. Non-SNP sites and sites with more than two alleles were removed. The 121 samples overlapping those phenotyped by van Boheemen, Atwater and Hodgins^14^ were retained (table S5), and sites with a minor allele frequency below 0.05 were removed, resulting in 1,142,278 SNPs for analysis. Genome-wide association studies (GWAS) were performed across 121 individuals from both North American (*n* = 43) and European (*n* = 78) ranges using EMMAX^75^, and incorporating an identity-by-state kinship matrix (generated in PLINK 1.9)^72^ to account for genetic structure among samples. The kinship matrix was computed using 790,209 SNPs which remained after pruning for linkage disequilibrium using a window size of 50kb, a step size of 5bp and an *r^2^* of 0.5. Candidate SNPs were identified using a conservative threshold of Bonferroni-corrected *p*-values < 0.05.

### Phenotypic analysis of herbarium specimens

We conducted a trait-based analysis of herbarium specimens found in the Global Biodiversity Information Facility database (gbif.org 2021). We compiled information from all *A. artemisiifolia* European herbarium specimens for which there was a digitized image of the individual in the database alongside corresponding metadata (location and collection date). The collection date spanned 1849 to 2020 (median 1975) and comprised 985 specimens. We determined the stage of flowering (no male inflorescence present, only immature male inflorescence present, mature male inflorescence present) for each image. The presence of fruit was also recorded. The male inflorescence was used as an indicator of flowering as these structures are more visually prominent than female flowers and the onset of male and female flowering is highly correlated^14^. Male florets consist of prominent spike-like racemes of male capitula, and are found at the terminus of the stem, whereas female florets are observed to be in inconspicuous cyme-like clusters and are arranged in groups at the axils of main and lateral stem leaves (fig. S4). The dates when the specimens were collected were converted to Julian day of the year. We conducted a generalized linear model with a binomial response and logit link (glm R). Both binary traits (presence of a mature male inflorescence; the presence of fruit) were included as response variables in two separate models. The significance of the effects were tested using the *Anova* function (Car package R)^76^ using type 3 tests. For both models, the predictors of latitude, day of the year, and collection year as well as all interactions were included. Non-significant interactions were removed in a stepwise fashion, starting with the highest order. Latitude of origin strongly correlates with flowering time in common garden experiments^14^ and we expected northern populations to evolve early flowering relative to the start of the growing season to match the shorter growing seasons in these areas. As a result, if local phenology has evolved to better match the local growing season we predicted a collection year by latitude interaction, as the relationship between latitude and the probability of flowering in wild collected accessions should change over time when controlling for the day of collection.

### Historic-modern genomic comparisons

To identify targets of recent selection, we compared historic and modern samples from twelve locations (five locations from North America and seven from Europe; table S12). Historic samples were grouped based on age of sample and proximity to a modern population. Analyses were performed in ANGSD^77^ using genotype likelihoods. For each population location we calculated pairwise nucleotide diversity (θ_π_) for historic and modern populations separately, and F_ST_ between historic and modern populations at each location. Statistics were calculated in non-overlapping 10kbp windows, and windowed θ_π_ values were normalized by dividing by the number of sites in each window. At each location, windows with θ_π_ more than two standard deviations below the mean in both historic and modern populations were excluded from the analysis. We identified putative selective sweeps in each population as windows with extreme shifts over time in allele frequency as well as extreme reductions in diversity (i.e. windows in the top one percent of both F_ST_ and θ_πH_/θ_πM_ distributions). To obtain further evidence for selective sweeps in these populations, we also performed genome scans of Fay and Wu’s H in each modern population. We first generated an ancestral consensus sequence in ANGSD (-*doFasta 2 -minMapQ 25 -minQ 20 -remove_bads 1 -uniqueOnly 1 -doCounts 1*) from alignments of *Ambrosia carduacea* and *Ambrosia chamissonis* to our *A. artemisiifolia* reference genome. We then used this ancestral sequence in calculating Fay and Wu’s *H* in 10kbp windows in ANGSD.

### Temporal allele frequency shifts in candidate loci

In order to track allele frequency shifts over time, we estimated contemporary and historical allele frequencies of the *ELF3* non-synonymous SNP and the haploblock hb-chr2, which are two candidate loci for recent selection in Europe. Both candidates showed evidence of local selection using spatial analysis of modern populations, as well as sweep signals in temporal comparisons of individual populations. These calculations were performed in geographic regions where this recent selection is believed to have occurred at both historic and contemporary timepoints. ANGSD^77^ (-*minMapQ 10 -minQ 5 -GL 2 -doMajorMinor 1 -doMaf 2 - doIBS 1 -doCounts 1 -doGlf 2*) was used to calculate the allele frequency of the early flowering *ELF3* allele (11:41517231) in 15 historic and eleven modern samples from within 200km of Berlin, whilst the frequency of hb-chr2 in Europe was ascertained using haploblock frequency estimates from across the European range (see below). To understand the magnitude of these allele frequency shifts relative to putatively neutral alleles elsewhere in the genome, we calculated a standardized measure of frequency change, *y_t_*, using estimates of historic, *p*_0_, and contemporary, *p_t_*, allele frequencies according to the equation:

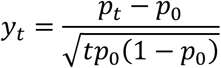

where *t* is the number of generations separating the frequency estimates (equivalent to the number of years due to ragweed’s annual lifecycle). As we show in supplementary text S3, the distribution of *y_t_* estimates under neutrality are predictable and roughly independent of the initial frequency of each neutral variant once the loci with low-frequency initial minor allele frequencies are filtered out. To further assess if selection was the likely cause of temporal changes of the *ELF3* and hb-chr2 variants, we estimated the distribution of *y_t_* estimates computed from 10,000 randomly-sampled SNPs that were located outside annotated genes and haploblocks and pruned for linkage disequilibrium using a window size of 50kb, a step size of 5bp and an *r^2^* of 0.5 in PLINK 1.9^72^. Prior to calculation of *y_t_*, sampled SNPs were then filtered for a minor allele frequency > 0.2 for hb-chr2 comparisons and MAF > 0.05 for *ELF3* comparisons (due to the low historic frequency of *ELF3* in historic Berlin populations), resulting in null distributions of between 6,913 and 8,303 SNPs. We then compared the distributions to the *y_t_* values of candidate adaptation loci to test whether candidate regions were more divergent than the putatively neutral distribution. As a point of comparison we repeated this analysis for hb-chr2 in North America and the *ELF3* allele in Quebec. As in Berlin, the *ELF3* allele is at high frequencies, but substantial temporal change was not expected as the populations were predicted to be closer to the equilibrium over the temporal sampling period in the native range. Samples within 200km of Quebec City (ten historic and eleven modern) were pooled at both timepoints. Allele frequency changes of the 10,000 randomly-sampled SNPs and the non-synonymous *ELF3* allele were assessed as above.

### Haploblock identification

To identify signatures of large, segregating haploblocks across the genome, we performed local windowed principal component analysis with Lostruct^43^. Using SNP data from 311 modern samples, we extracted the first ten multidimensional scaling (MDS) coordinates across each chromosome in windows of 100 SNPs. These MDS coordinates were then plotted along each scaffold to observe regions of local structure, indicative of segregating haploblocks. We focused on outlier MDS signals that overlapped parallel outlier windows for both XtX and at least one environmental variable, and also showed well-defined boundaries indicative of chromosomal inversions. We tested for additional evidence of inversions using PCA of MDS outlier regions and kmeans clustering in R^74^ to identify regions containing three distinct clusters representing heterozygotes and two homozygotes. Additionally, we assessed heterozygosity from genotype data in each haploblock region and in each modern sample, and measured linkage disequilibrium (the second highest *r^2^* value in 0.5Mbp windows) across each scaffold bearing a haploblock for all modern samples and for modern samples homozygous for the more common haploblock genotype using scripts from Todesco *et al.^26^*.

### Haploblock frequency changes over time and space

For fifteen candidate inversions, a local PCA of each region and kmeans clustering was then repeated in PCAngsd^78^, so as to allow genotype estimation of these haploblocks in 305 historic samples alongside the 311 modern samples. For this local PCA we used only the chromosomal regions already defined as haploblocks in order to obtain population wide clustering for both historic and modern datasets, which we then used to infer haploblock genotypes. We also conducted a PCA on 10,000 SNPs randomly-sampled from the 311 modern genomes that were located outside annotated genes and haploblocks, and pruned for linkage disequilibrium using a window size of 50 kb, a step size of 5 bp and an *r^2^* of 0.5 in PLINK^72^. Following this, we used generalized linear models (glm R) to assess how haplotype frequency (binomial response) changed over time and space. A count of each haplotype at a geographic location and year was the binomial response variable and time period (historic or modern), range (North America or Europe), latitude, and all interactions between these three main effects were used as predictors. Non-significant interactions were removed in a stepwise fashion, starting with the highest order. PC1 from the PCA of 10,000 randomly-sampled SNPs was included as a covariate to control for the effects of population structure on haplotype frequency. We tested the significance of the effects in our model using the Anova function (Car package R)^76^ with type 3 tests. Significant differences among groups for means or slopes were tested with the emmeans package using an FDR correction^79^. To determine if the classification of samples into modern or historic timepoints influenced our results we ran a second set of generalized linear models examining haplotype frequency as a function of collection year, range (North America or Europe), latitude, and all interactions between these three main effects as well as PC1, using the same approach as above. For interactions involving two continuous variables (i.e., latitude and year) we tested if the slope estimates of one variable were significant at specific values of the other using the package emmeans. This allowed us to estimate when and where the haplotype frequencies were changing. The results from both approaches (time as two categories or time as continuous) provided qualitatively similar patterns.

We estimated the relative strength of selection on haploblocks along the latitudinal clines in modern North American and European populations using slopes from logistic regressions (see supplementary text S2). Specifically, we used generalized linear models to estimate the slopes of the regression for each range and time point (modern or historic) combination (group). A count of each haplotype at a geographic location and collection year was the binomial response variable and time period (historic or modern), range (North America or Europe), latitude, and all interactions between these three main effects were used as predictors. All interactions were retained in the model and slopes and their confidence intervals estimated for each group using the function emtrends (emmeans package R^79^; table S14). PC1 was included as a covariate to control for the effects of population structure on haplotype frequency. We expected the slopes to be shallower in the historic versus the modern European group, but similar across timepoints in North America. To test this, we used a t-test and compared slopes for modern and historic timepoints in each range. We also expected that the magnitude of change in the slope over time would be the greatest in haploblocks showing the largest estimates of selection in Europe (table S16). We estimated the relative strength of selection for the modern European range for each haploblock and tested if the absolute change in slope for each haploblock was correlated with this estimate. We also examined if there was a correlation in the relative strength of selection for modern North American and European haploblocks, which would indicate parallel selection along the cline in each range.

We compared our slope estimates of the haploblocks to the genome wide distribution in each range using 10,000 randomly selected SNPs outside of genes and haploblocks. We did this to determine if our haploblocks showed stronger latitudinal patterns than the majority of SNPs, in one or both ranges, which may be indicative of spatially varying selection. For the modern samples in each range (North America or Europe), we fit a generalized linear model with latitude as the only predictor. We did this for each null SNP and each haploblock that was statistically associated with latitude.

### Recombination rates in haploblocks

The haploblocks show multiple genomic signatures of reduced recombination. To confirm this we analyzed recombination rates in genetic maps. Further if the haploblocks were caused by global reductions in recombination rate (e.g., the region was found in an area with generally low recombination such as a centromere), all maps should show reduced recombination rates. However, if inversions were the cause, recombination would only be suppressed in genotypes heterozygous for the inversion, while homozygous individuals would not show suppressed recombination. To determine if there were genotype-specific reductions in recombination rate in the haploblocks, which would be consistent with inversions, we made use of three previously generated genetic maps^40^. Markers were generated using genotype by sequencing and alignments to the haplotype 1 of our diploid reference genome. Details of the sequencing, alignments and variant calling can be found in Prapas *et al.^40^*. We developed sex-specific genetic maps (i.e., maps for the maternal and paternal parent) using Lep-MAP3^80^ for each chromosome of interest and in each mapping population (an F1 mapping population and two F2 mapping populations). Multiple maps were constructed since the haploblocks may have been segregating in different frequencies in the parents of the mapping populations derived from outcrossing. For the recombination rates, linkage map construction was constrained by the physical order of the markers along each scaffold of interest. Genetic distance (cM) was plotted against physical position along the chromosome for each map and the intervals of the QTL and the boundaries of the haploblocks were visualized and inspected for reduced recombination compared to the rest of the scaffold. We also used the genetic map (pink family) to confirm the OmniC scaffolding, and assess the accuracy of the manual correction of chromosome 18 in both assemblies (fig. S16).

## ACKNOWLEDGEMENTS

We thank Greg Owens & Michael Whitlock for discussions, and Samuel Yeaman and Sarah Otto for feedback on the manuscript. We are grateful to François Bretagnolle, Myriam Gaudeul, Heinz Mueller-Schaerer, Gerhard Karrer, and Bruno Chauvel for their assistance in obtaining many of the samples upon which this study is based, and Marie Brunier, Fátima Sánchez Barreiro, Yohann Roy, Luna Forcioli, Jacqueline Y. Lee for assistance during lab work. Some sequencing services were provided by the Norwegian Sequencing Centre, a national technology platform hosted by the University of Oslo. Some sequencing was performed by the NTNU Genomics Core Facility. Genome scaffolding was performed by Dovetail Genomics. Some analyses were performed on resources provided by Sigma2, MASSIVE M3 and ComputeCanada high performance computing platforms. We kindly thank the curators from the following herbaria for allowing us to destructively sample their precious collections: B, BR, BRNU, C, FI, G, GH, GOET, GZU, HBG, I, IASI, JE, L, LD, LY, MARS, MASS, MO, MPU, NEBC, NEU, NY, P, PH, PR, PRA, PRC, QFA, S, STU, TRH, UPS, US, W, WU.

## AUTHOR CONTRIBUTIONS

**Paul Battlay**: Software, Formal analysis, Investigation, Data curation, Writing - Original draft, Writing - Review and editing, Visualization **Jonathan Wilson**: Software, Formal analysis, Investigation, Writing - Review and editing, Visualization **Vanessa C. Bieker**: Software, Investigation, Data curation **Chris Lee**: Investigation, Resources **Diana Prapas**: Software, Formal analysis **Bent Petersen**: Software **Sam Craig**: Investigation **Lotte van Boheemen**: Investigation, Resources **Romain Scalone**: Resources **Nissanka P. de Silva**: Software, Visualization **Amit Sharma**: Investigation, Resources **Bojan Konstantinović**: Investigation, Resources **Kristin A. Nurkowski**: Investigation, Resources **Loren Rieseberg**: Writing - Review and editing **Tim Connallon**: Methodology, Formal analysis, Investigation, Writing – Original draft, Writing - Review and editing **Michael D. Martin**: Conceptualization, Methodology, Resources, Writing - Review and editing, Supervision, Project administration, Funding acquisition **Kathryn A. Hodgins**: Conceptualization, Methodology, Software, Formal analysis, Investigation, Resources, Writing - Original draft, Writing - Review and editing, Supervision, Project administration, Funding acquisition, Visualization

## DATA AND CODE AVAILABILITY

Sequences used in reference genome assembly and annotation are available from NCBI under BioProject ID PRJNA819156. Reference genome FASTA and annotation GFF files are available from FigShare (DOI TBA). Individual sample resequencing data are available from ENA under BioProject IDs PRJEB48563, PRJNA339123 and PRJEB34825.

## SUPPLEMENTARY FIGURES

**Figure S1.**
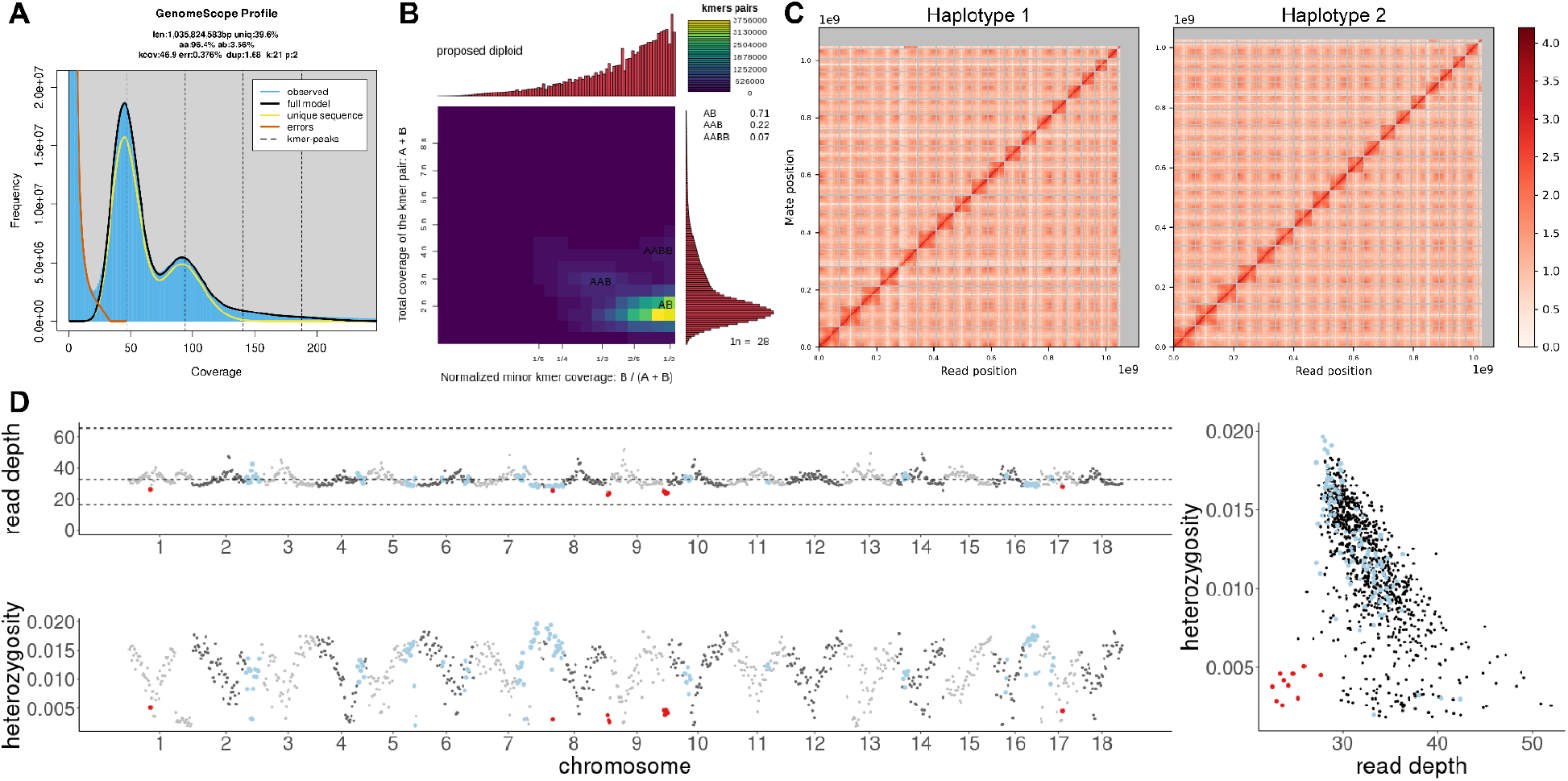
*Ambrosia artemisiifolia* genome assembly quality. **A**. The kmer coverage and model fit from GenomeScope 2.0 for 21mers from the PacBio HiFi and OmniC Illumina reads. **B**. A smudgeplot used to estimate ploidy from heterozygous kmer pairs using 21mers from the HiFi reads. **C**. Link density histograms, identified by proximity ligation sequencing for haplotype 1 and haplotype 2. The *x* and *y* axes show mapping positions of the first and second read in read pairs. **D**. Read depth and heterozygosity in 1Mbp windows for Illumina reads used in the reference genome assembly mapped to the final version of the reference genome. Windows in the bottom 10% of read depth and heterozygosity values are indicated in red; windows overlapping haploblocks are indicated in pale blue.

**Figure S2.**
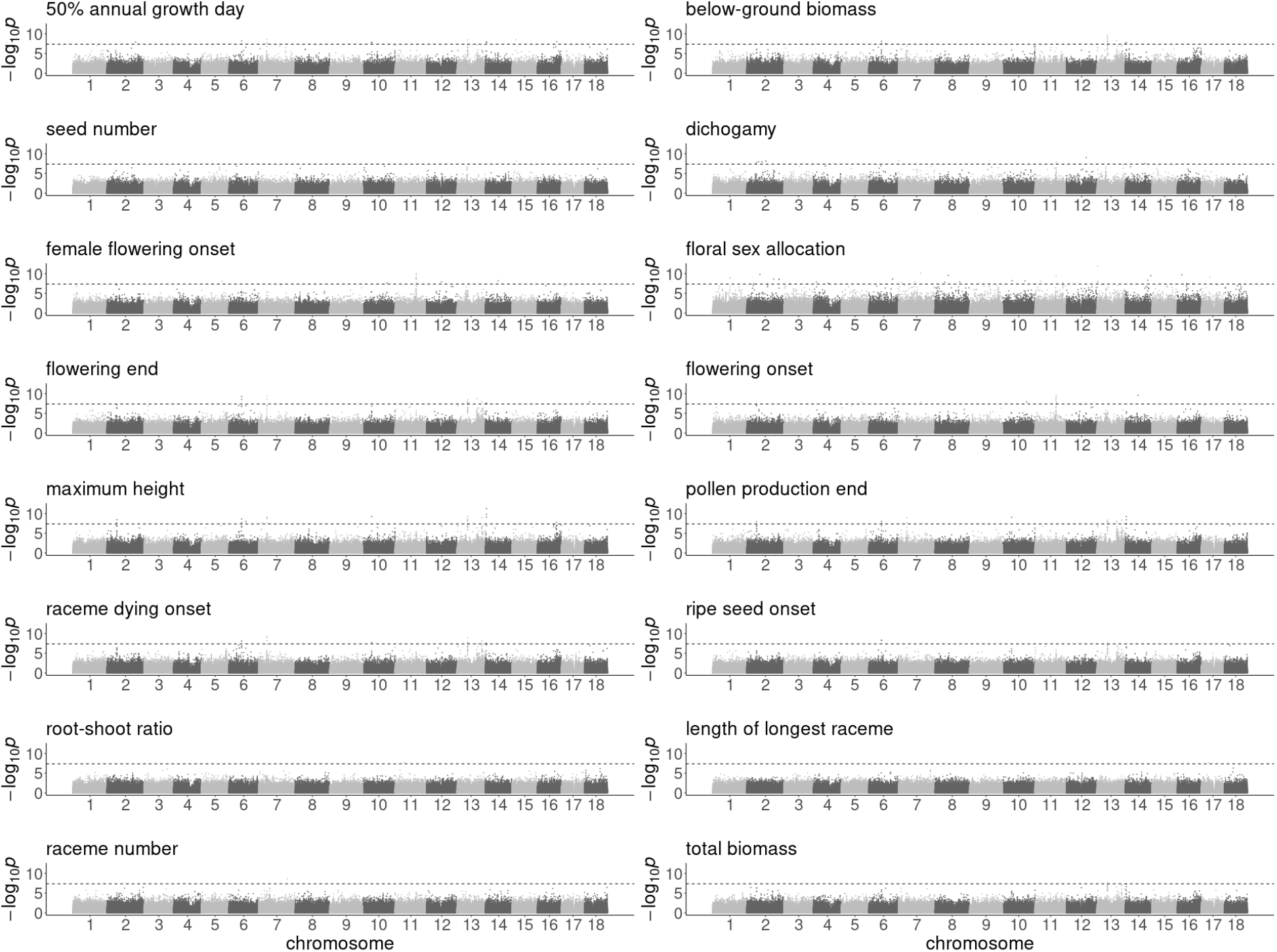
Genome-wide association study results for phenotypes with significant SNPs. Solid lines indicate a bonferroni-corrected significance threshold of 0.05.

**Figure S3.**
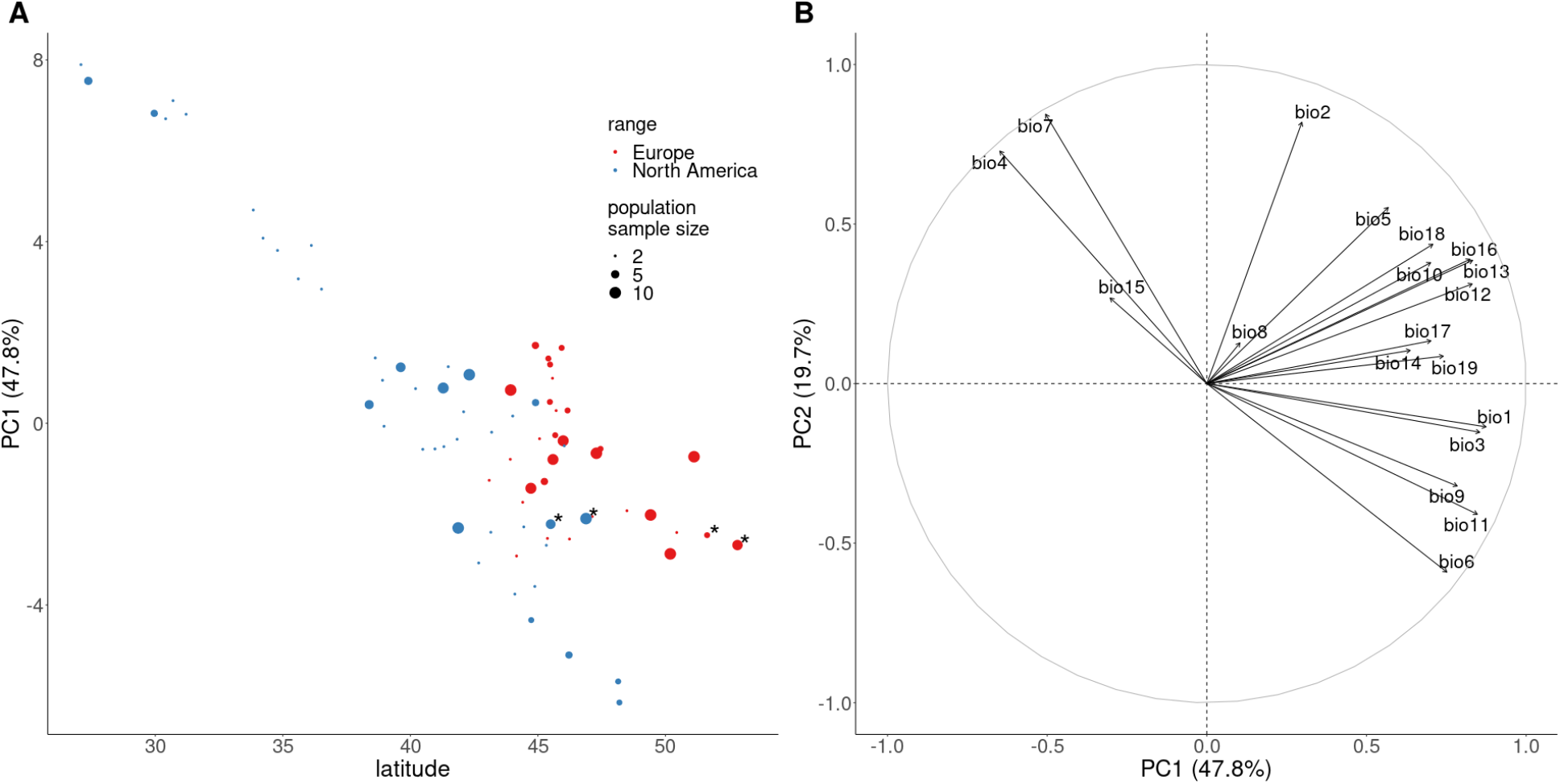
Environmental comparison of North American and European *A. artemisiifolia* ranges. **A**. The first principle component of 19 bioclimatic variables against latitude for modern ragweed populations in North America (red) and Europe (blue), with asterisks identifying populations with high *ELF3* allele frequencies. **B**. A variable correlation plot for 19 bioclimatic variables.

**Figure S4.**
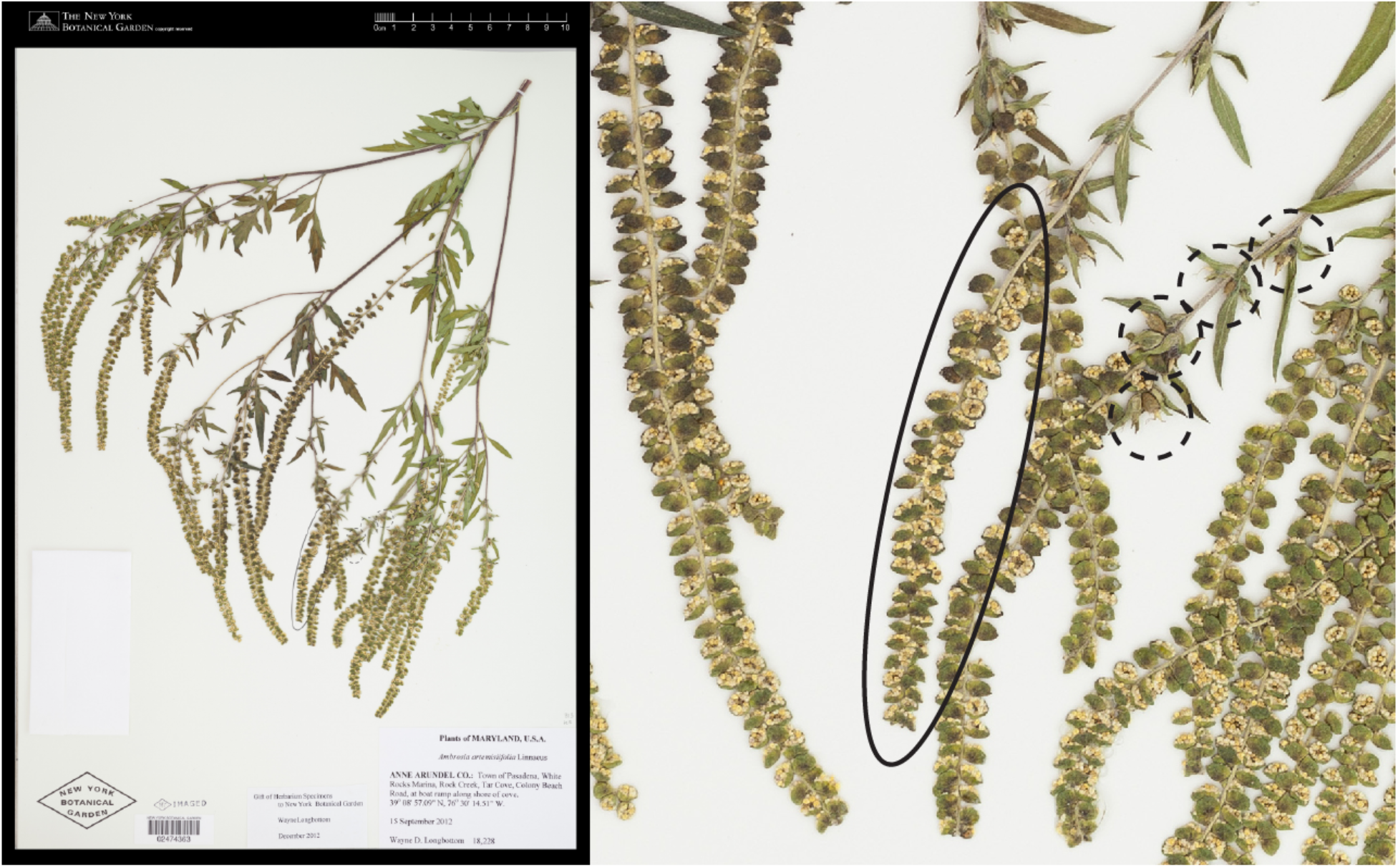
An example herbarium specimen of *Ambrosia artemisiifolia* (left) and a detail (right) showing the mature male inflorescence (solid line) and seeds (dashed lines).

**Figure S5.**
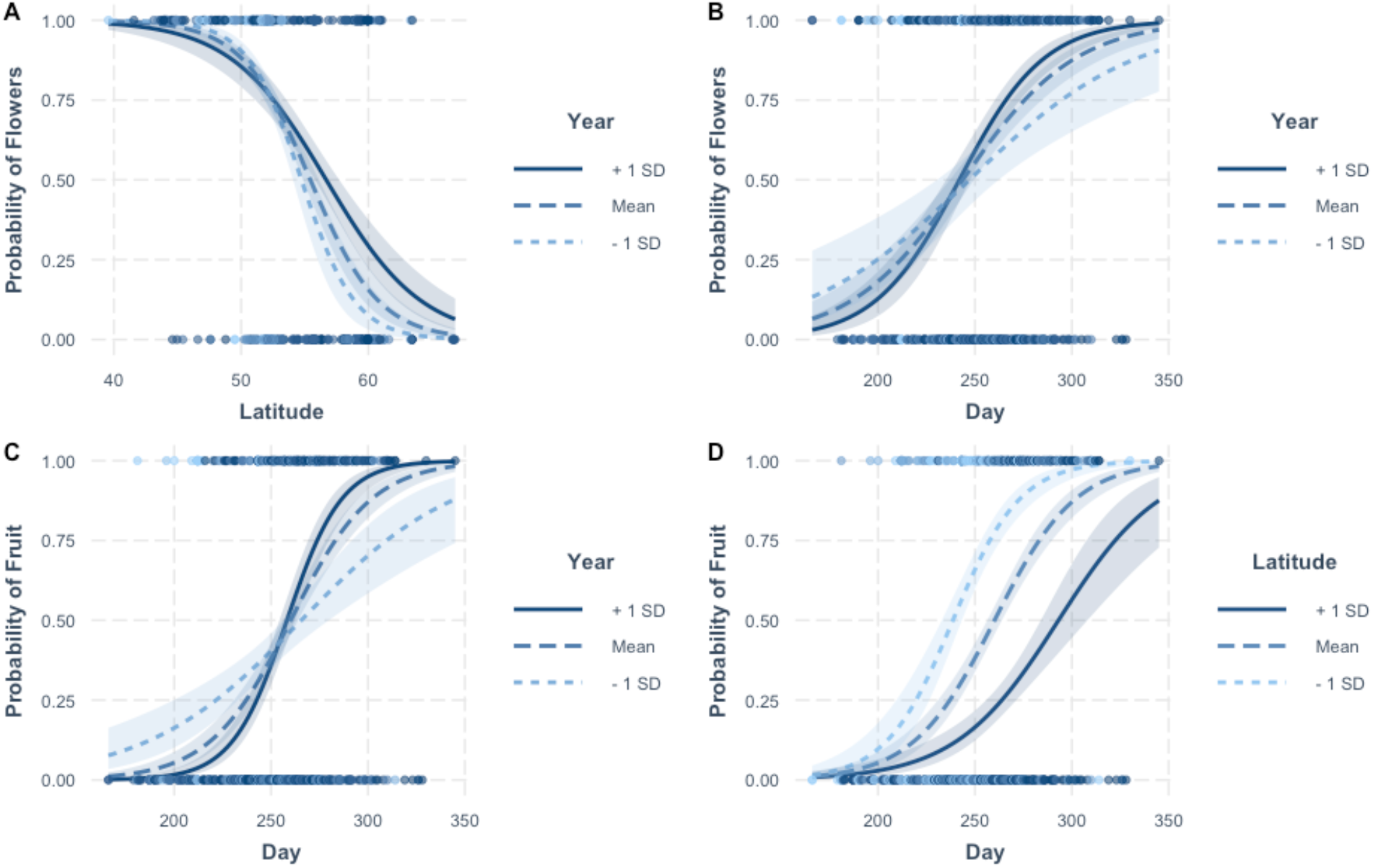
Interaction plots illustrating the results of generalized linear models examining the presence of mature male inflorescences (probability of flowers) or mature fruit (probability of fruit) in herbarium specimens of *A. artemisiifolia* in Europe as a function of collection day (Day), latitude of origin (Latitude) and collection year (Year). The predicted probability of observing flowers is plotted as a function of latitude (**A**), or collection day (**B**) for different collection years (mean collection year +/− 1 SD). The predicted probability of observing fruit is plotted against collection day for different collection years (mean collection year +/− 1 SD; **C**) or latitudes (mean collection latitude +/− 1 SD; **D**). Confidence intervals for the predictions are shown as are the raw data.

**Figure S6.**
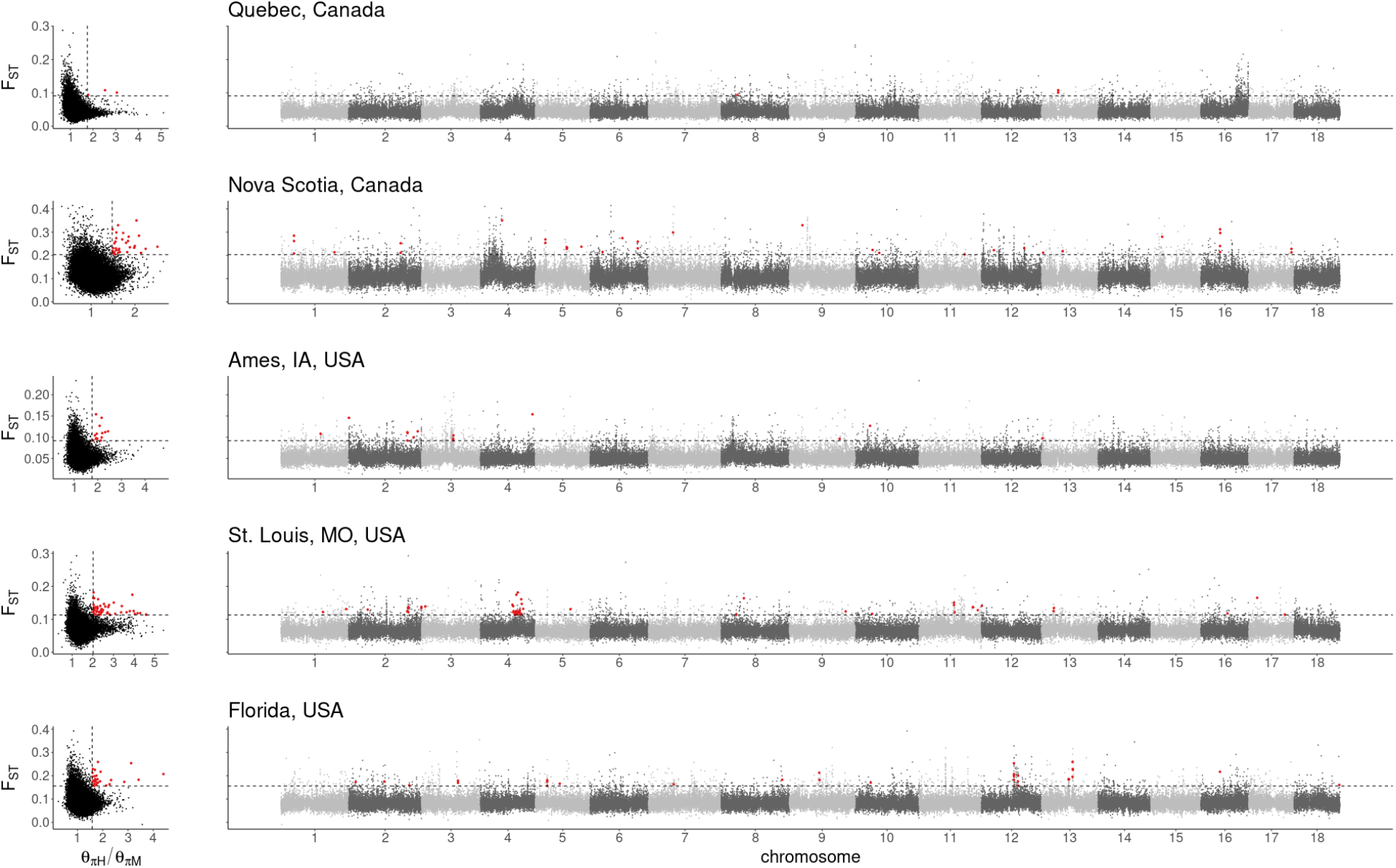
Distributions of F_ST_ and θ_πH_/θ_πM_ between historic and modern samples from North American populations, and F_ST_ against genomic location. Red points indicate putative selective sweep windows, which are in top one percent of per-window F_ST_ and θ_πH_/θ_πM_ (dashed lines).

**Figure S7.**
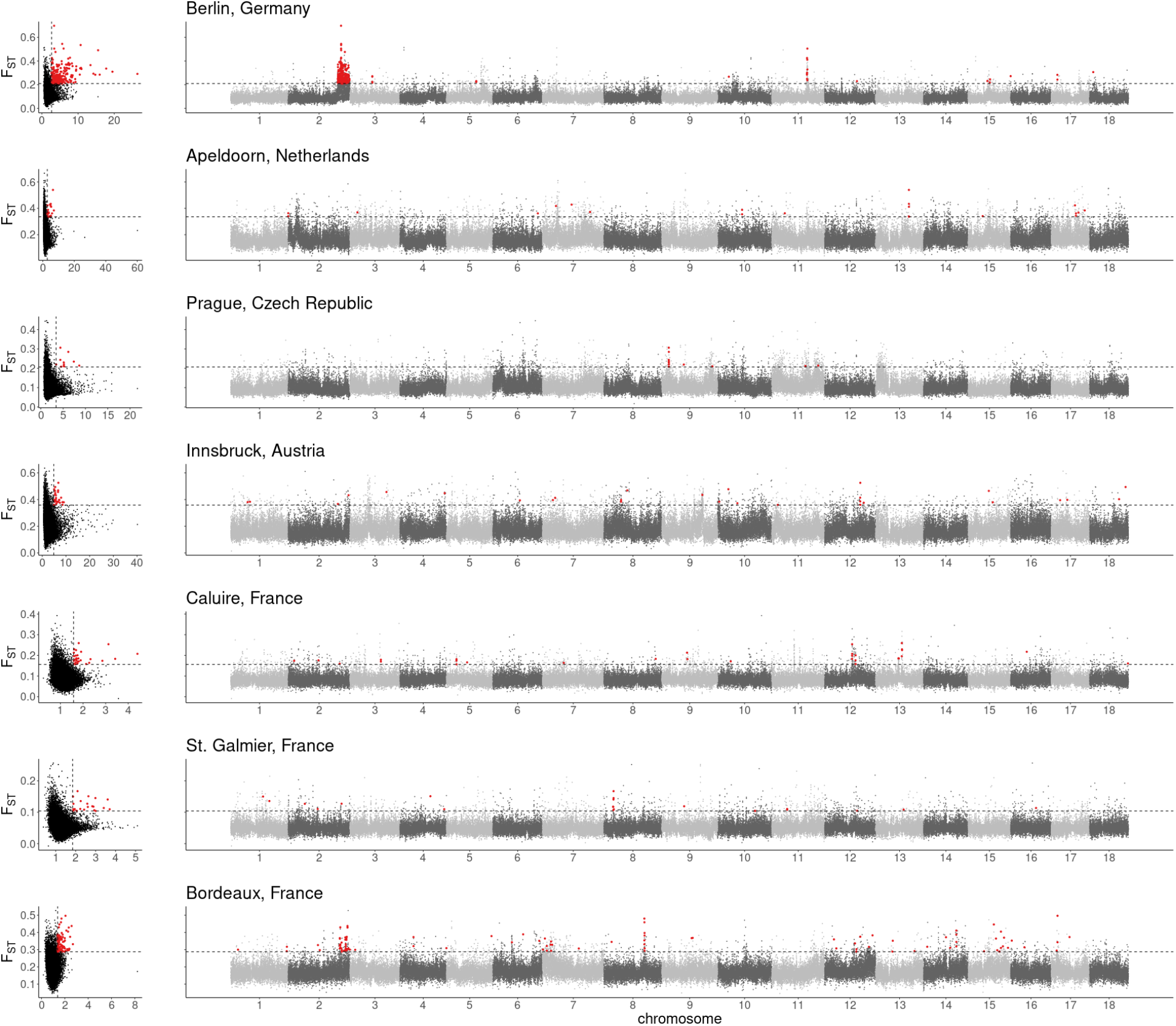
Distributions of F_ST_ and θ_πH_/θ_πM_ between historic and modern samples from European populations, and F_ST_ against genomic location. Red points indicate putative selective sweep windows, which are in top one percent of per-window F_ST_ and θ_πH_/θ_πM_ (dashed lines).

**Figure S8.**
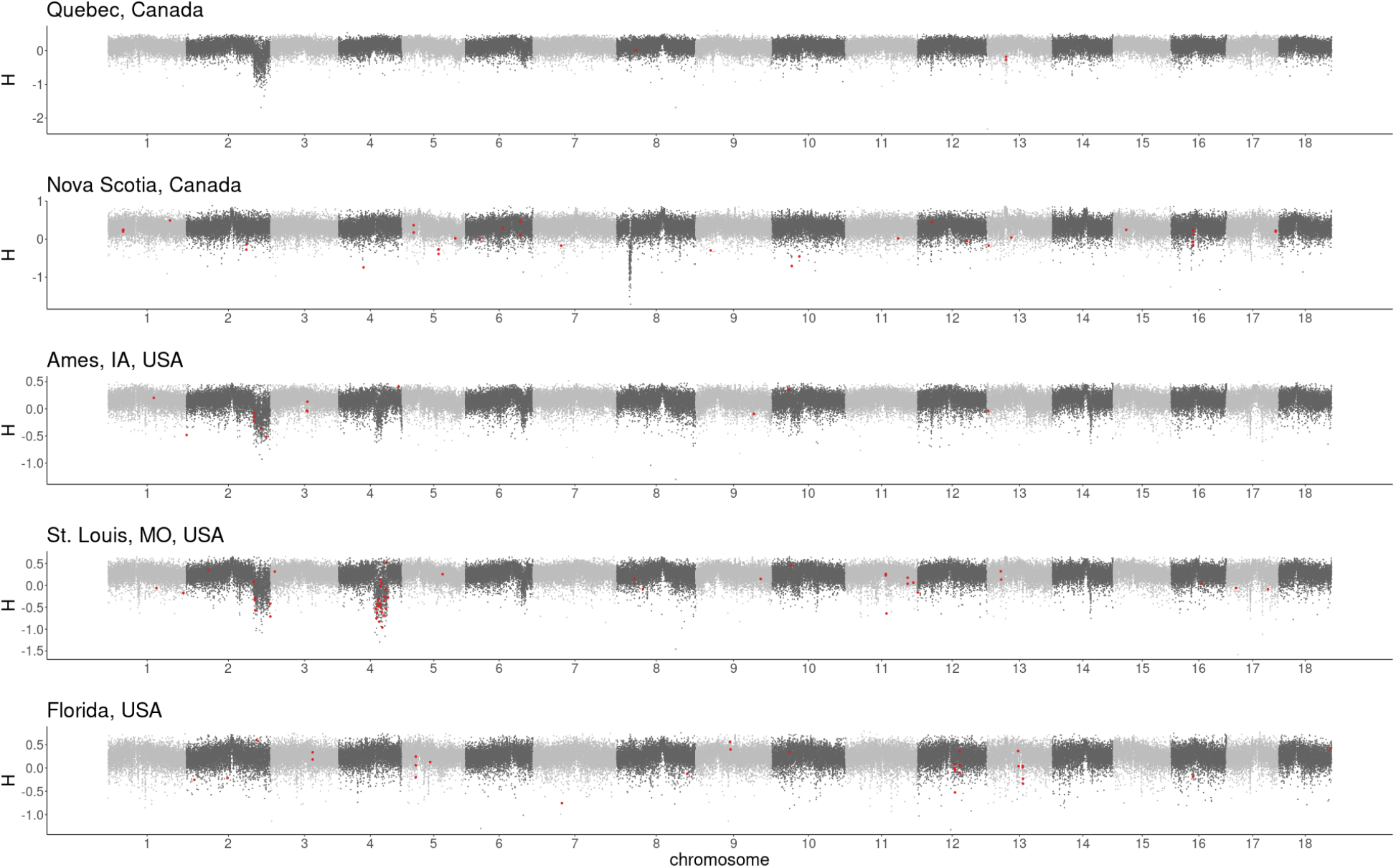
Fay and Wu’s *H* against genomic location for modern North American populations. Red points indicate putative selective sweep windows from historic-modern comparisons.

**Figure S9.**
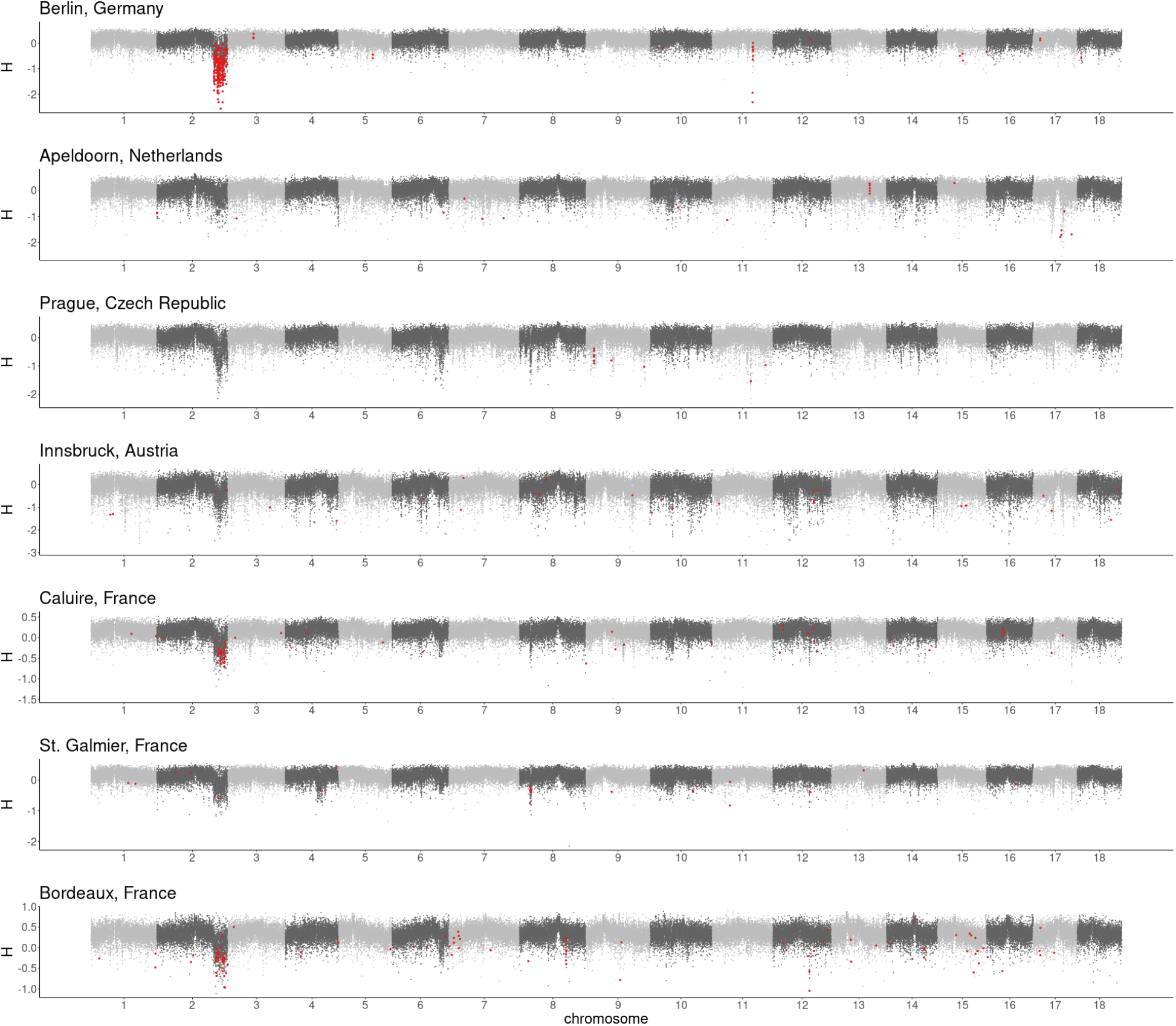
Fay and Wu’s *H* against genomic location for modern European populations. Red points indicate putative selective sweep windows from historic-modern comparisons.

**Figure S10.**
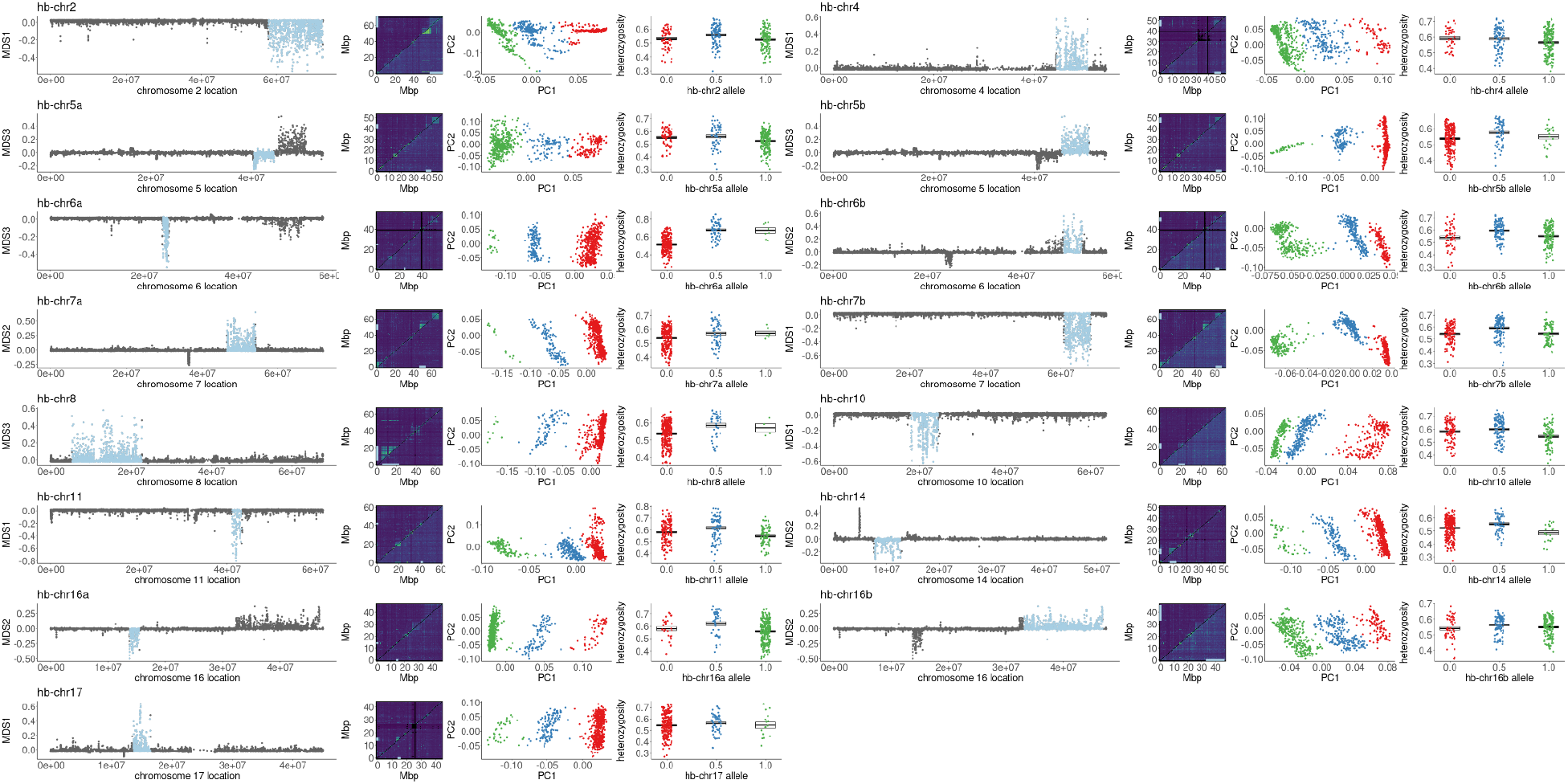
Haploblocks display extreme, divergent, local population structure (pale blue regions; first column). Haploblock regions (indicated by pale blue lines; second column) correspond to blocks of linkage disequilibrium (second highest *r^2^* in 0.5Mb windows) apparent using all modern samples (top triangle) but often reduced or absent using only samples homozygous for the more common haploblock genotype (bottom triangle). Haploblock genotypes were assigned by kmeans clustering (colours; third column) using the first two principal components of genetic variation across haploblock regions. Heterozygous haploblock genotypes show elevated mean per-site heterozygosity (fourth column; boxes denote mean and SEM).

**Figure S11.**
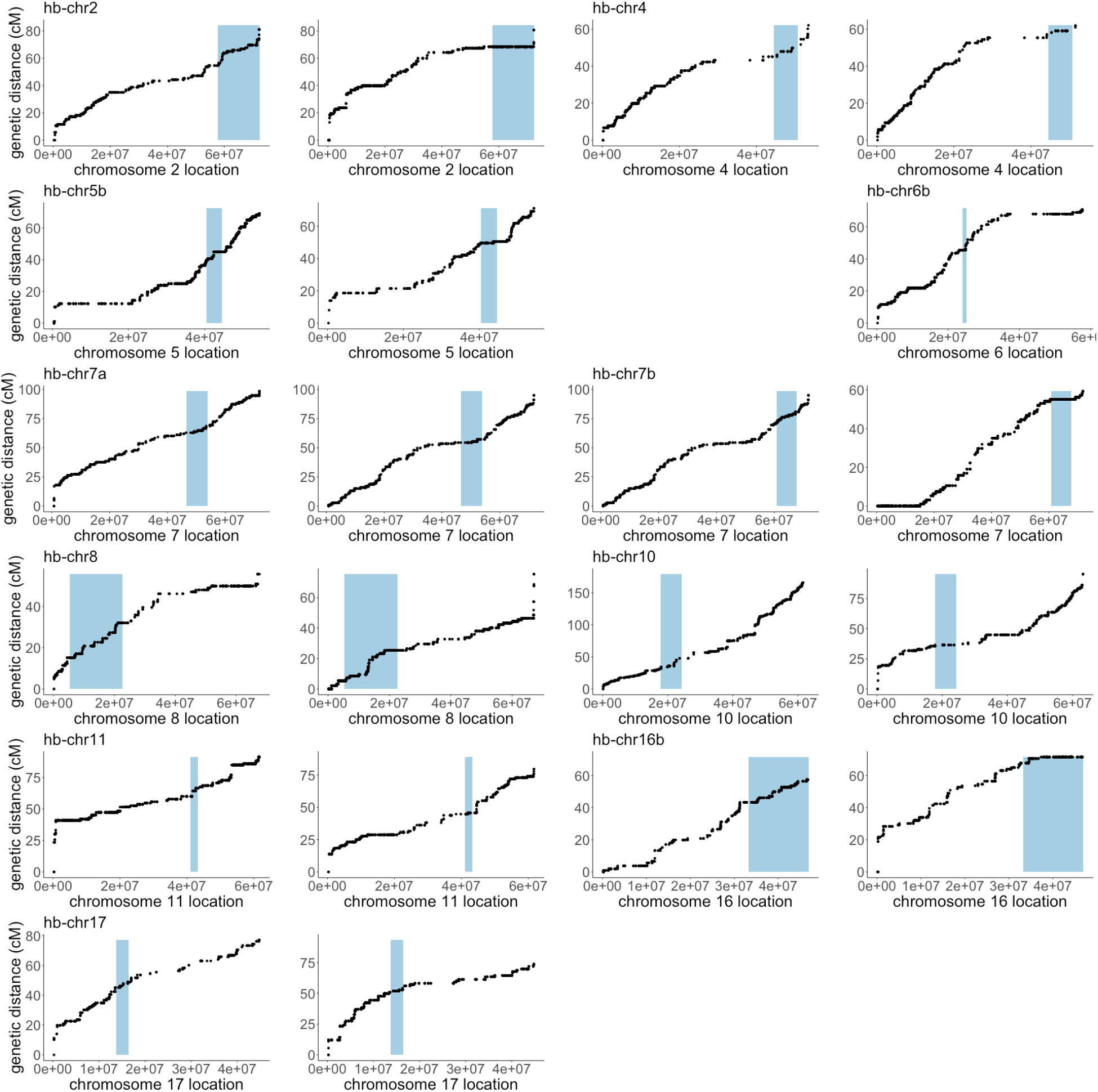
Evidence of genotype-specific reductions in recombination in haploblock regions. Genetic distance (cM) against physical distance (bp) along a portion of each scaffold is shown. Haploblock regions are shown in pale blue. Example maps are displayed showing both low recombination rates (left) and high recombination rates (right) for each haploblock.

**Figure S12.**
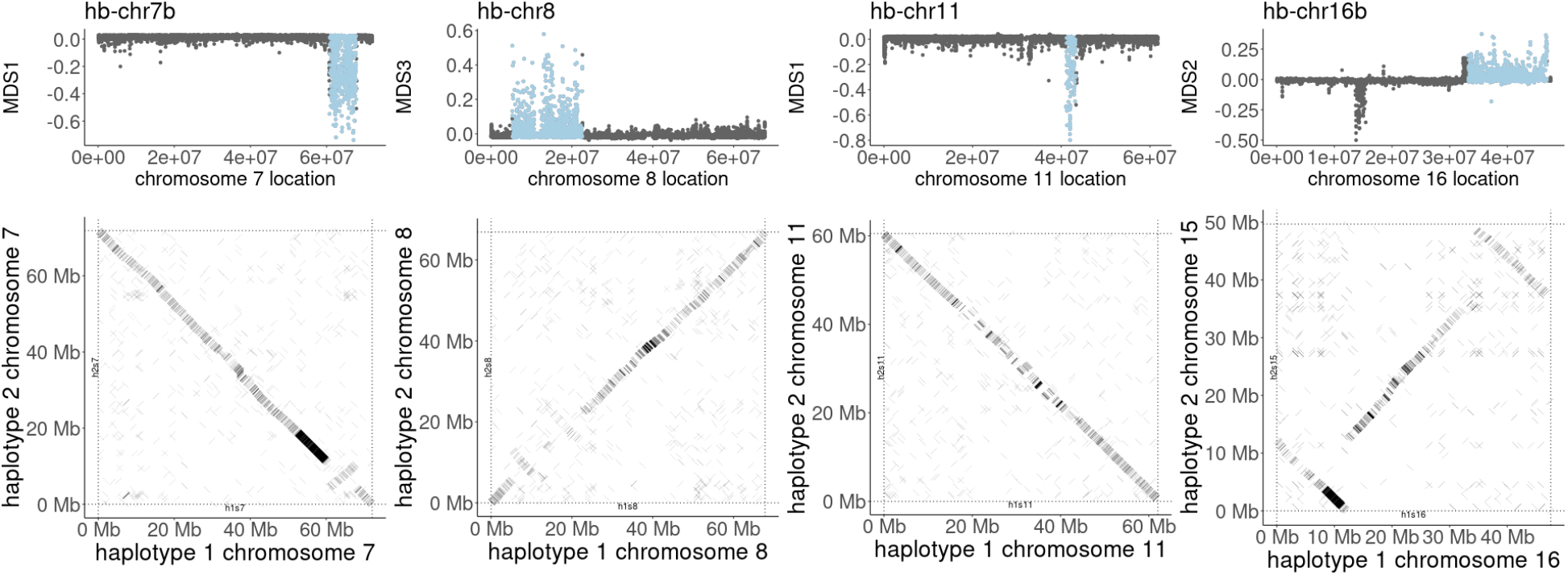
Population-genomic signatures of inversions (top row; from fig. S9) that correspond to inversion polymorphisms segregating in our diploid reference genome (bottom row; from fig. 1B).

**Figure S13.**
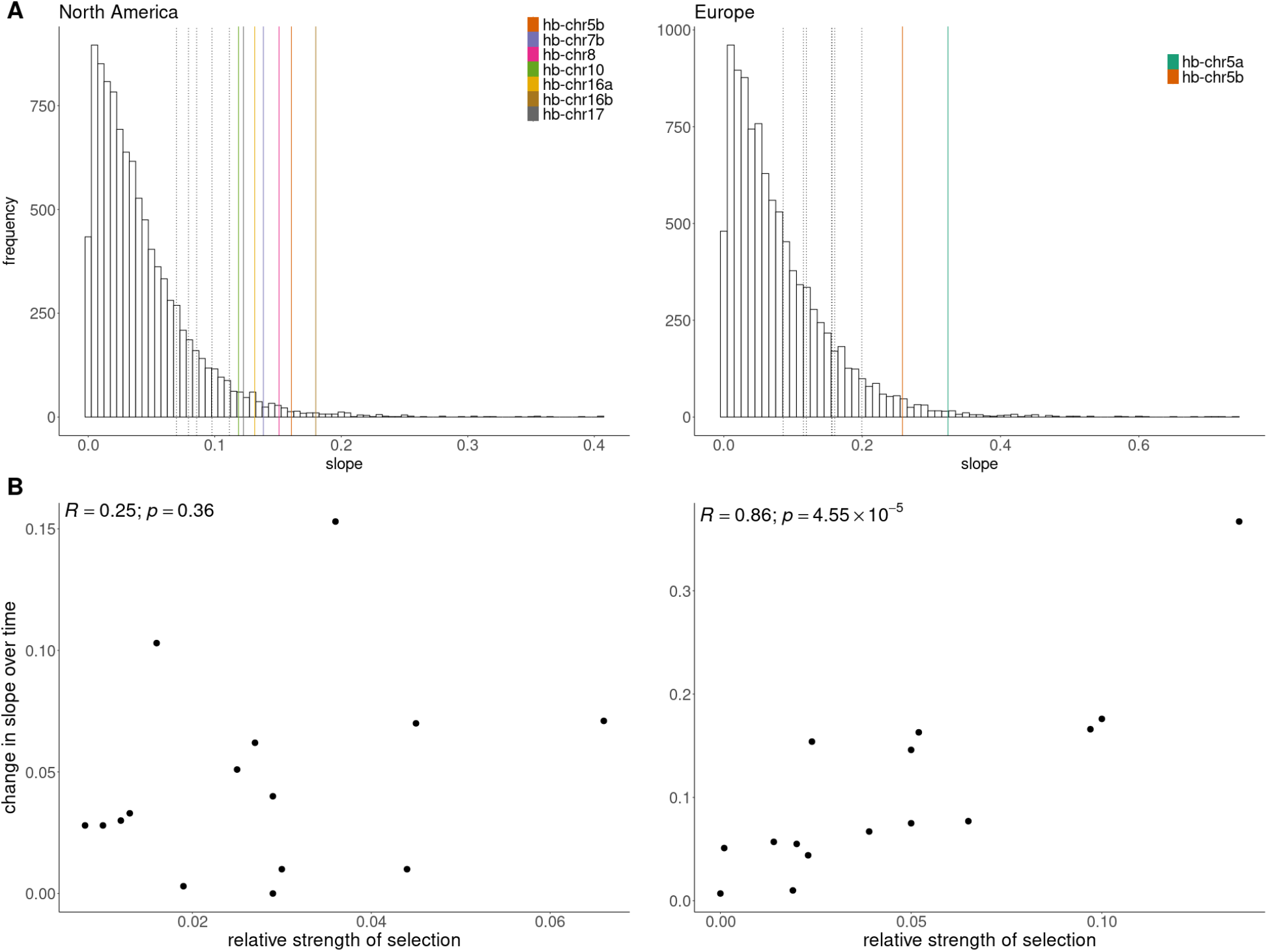
**A**. The distribution of slope estimates from generalized linear models of population allele counts against latitude for 10,000 randomly selected SNPs in each range. The vertical lines show the slope estimates for haploblocks with statistical associations with latitude in one range (table S12). Solid lines represent estimates in the 5% tail of each distribution while dotted lines fall below that cut-off. **B**. The change in the slope of the relationship between latitude and haplotype frequency (see table S15) between historic and modern samples compared to the estimate of selection along the latitudinal cline for the 15 haploblocks (estimated from modern data in each range). A strong relationship was detected in the invasive European, but not the native North American range.

**Figure S14.**
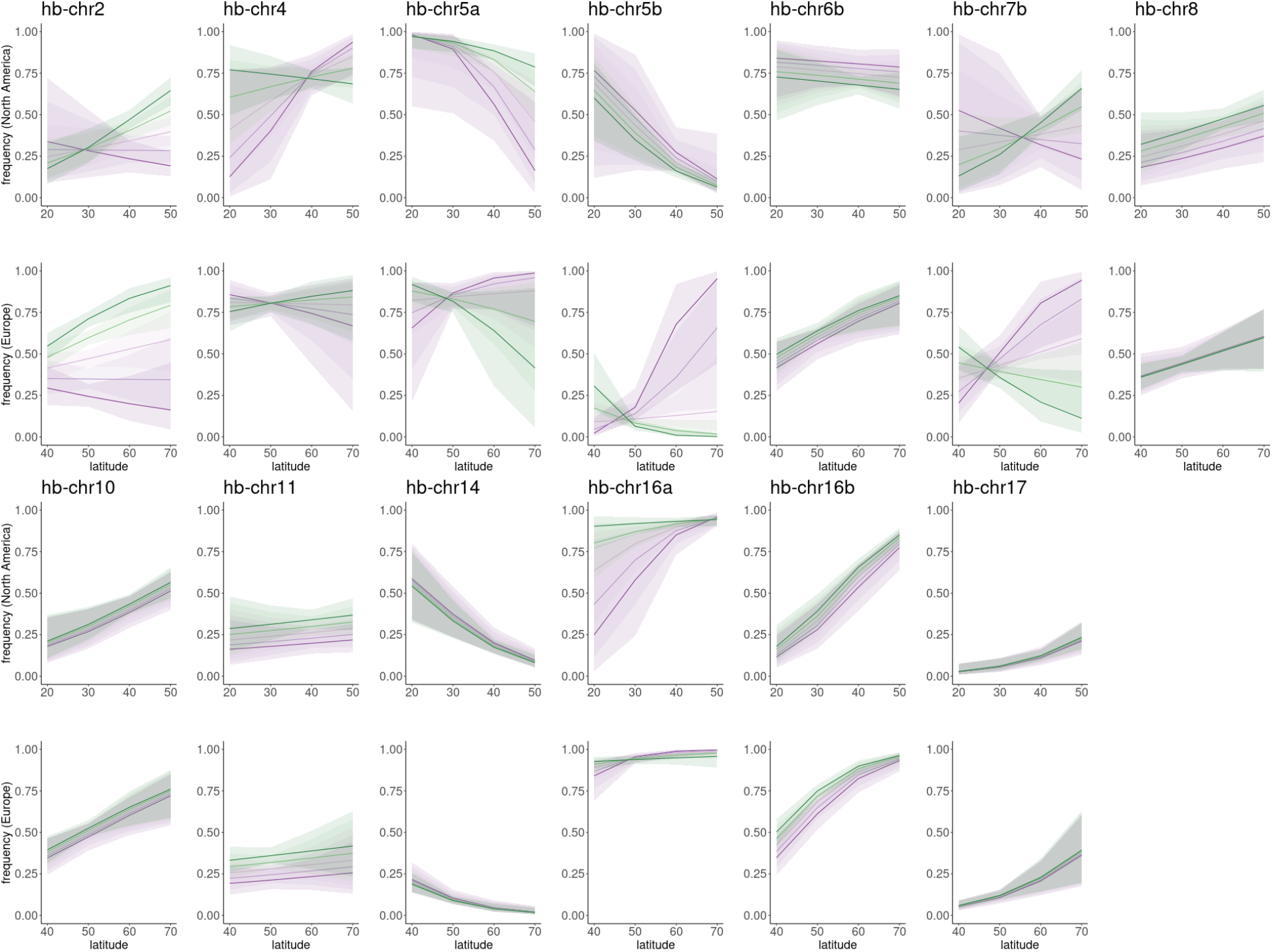
Logistic regression models with 95% CI ribbons (see table S12-S17 for model details) of haploblock frequency (allele 1) against latitude for each haploblock that shows a significant latitude, time or range effect, or significant interactions between these effects, across five time bins ranging from most historic (purple) to most modern (green).

**Figure S15.**
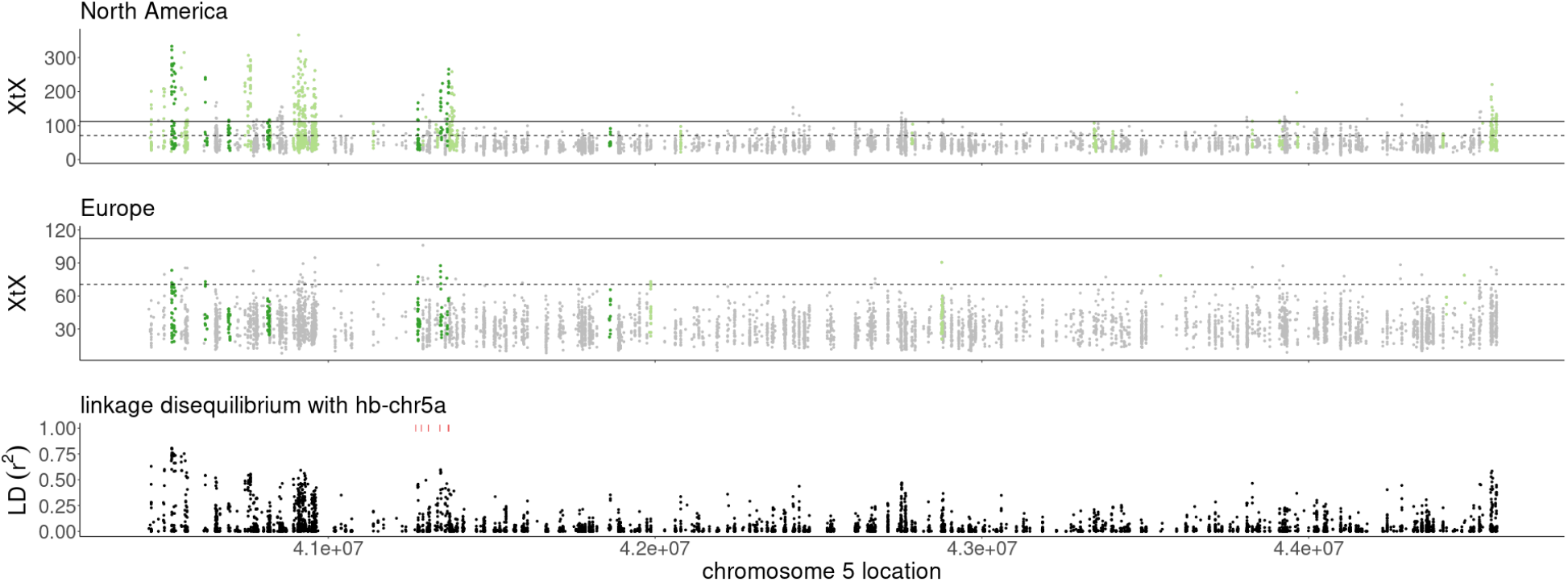
A detailed analysis of the hb-chr5 region (5:40446189-44576095). XtX and XtX-EAA outlier windows zoomed from fig. 2C, and linkage disequilibrium (*r^2^*) between SNPs in the region and hb-chr5a haploblock genotype. A cluster of six pectate lyase genes, consisting of the top BLAST hit for *Amba1* and closely-related paralogues, are indicated in red above the linkage disequilibrium plot.

**Figure S16.**
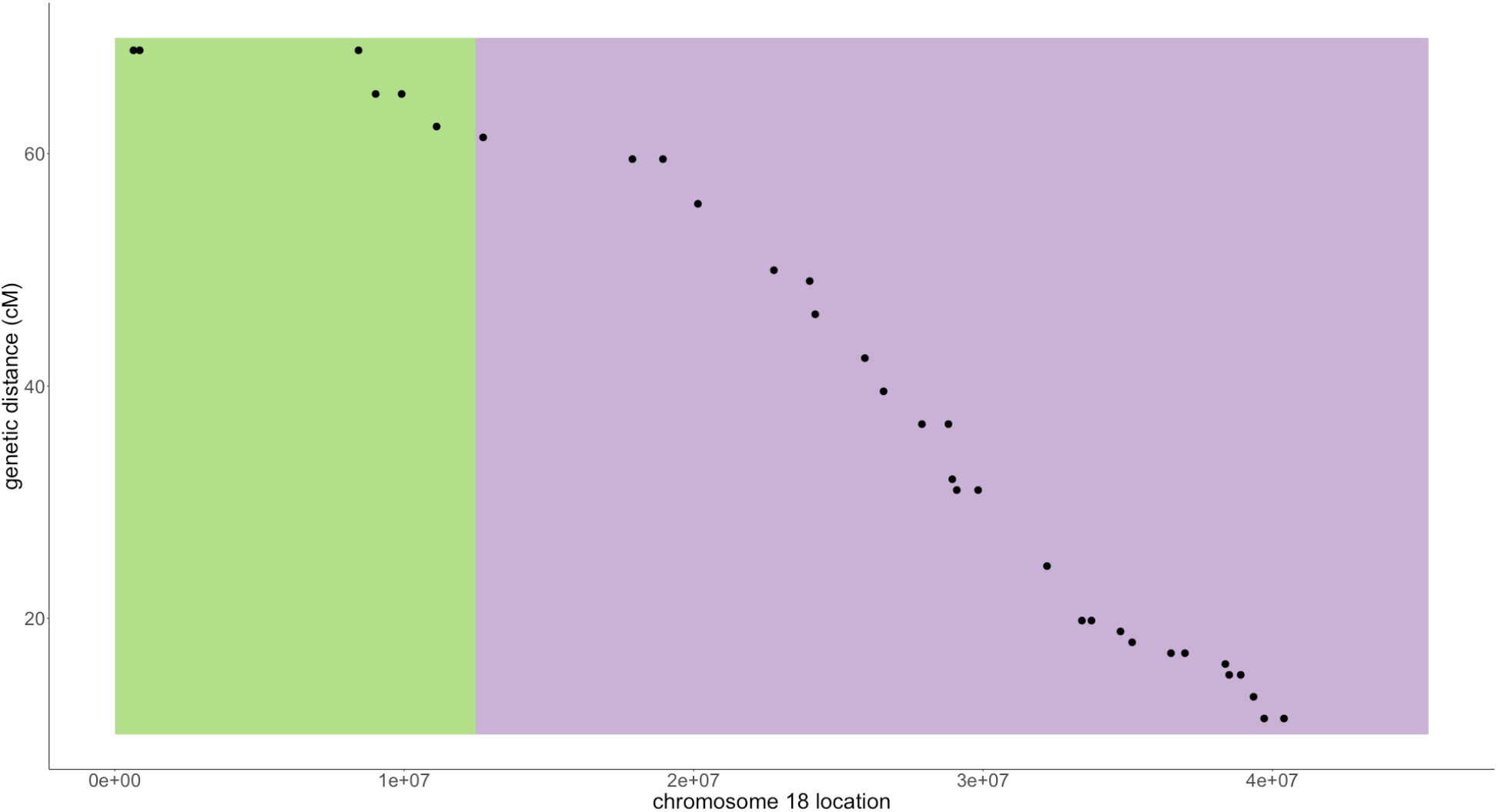
Genetic markers support the combination of two scaffolds (green and purple) to assemble haplotype 1 chromosome 18.

**Figure S17.**
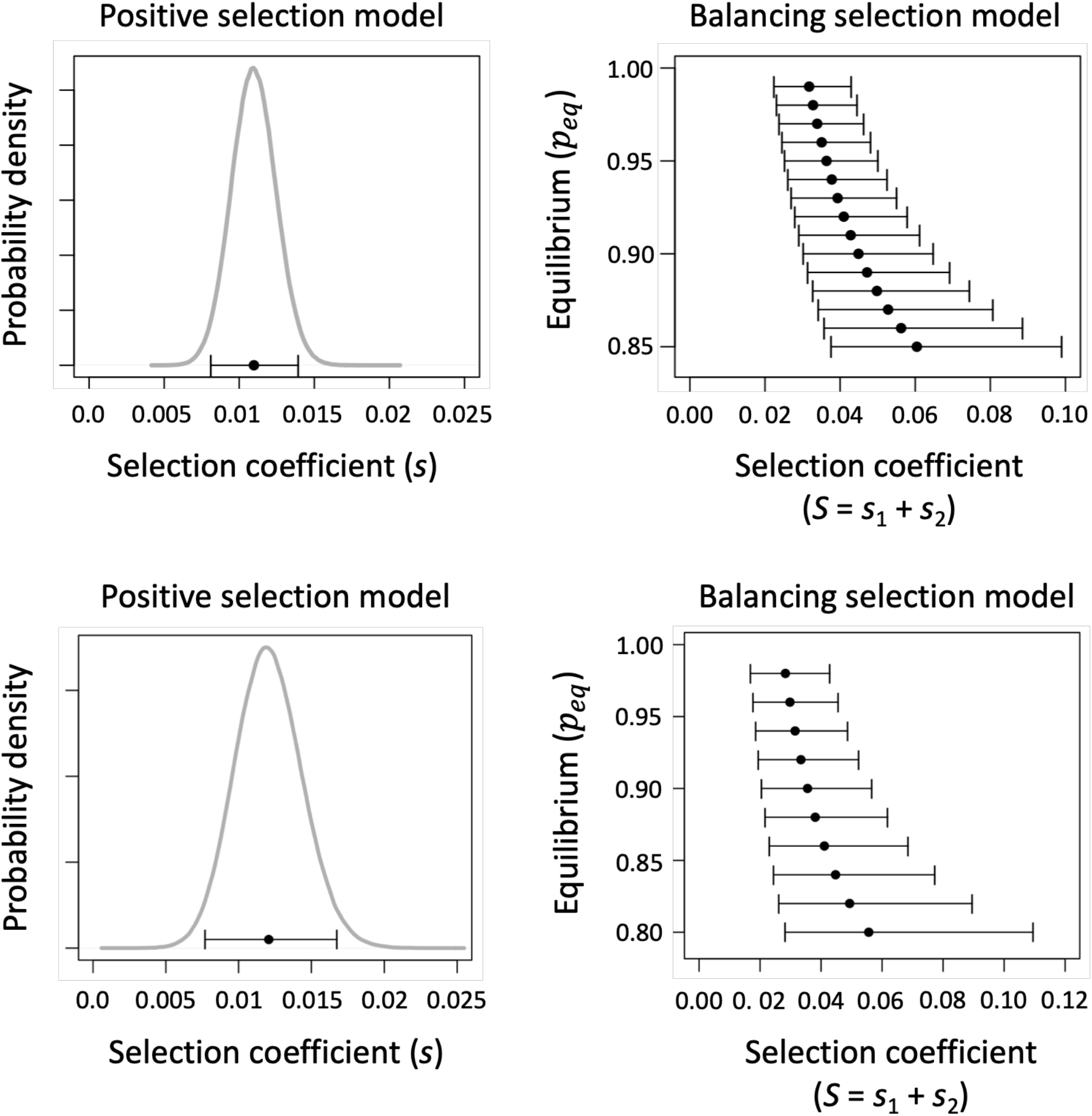
Selection coefficients consistent with observed frequency changes of the hb-chr2 haplotype, under models of positive selection (left) and balancing selection (right). The left-hand panel shows the distribution of *s* values (gray) and the 95% CI for *s* (in black: parallel to the *x*-axis), consistent with temporal change in the hb-chr2 haploblock. The distribution of s is based on simulations of 10^6^ initial and final frequencies of the haploblock that are consistent with the estimated frequencies and error in the estimates. Eq. (S1) was used to calculate a value of *s* for each set of initial and final frequencies during the time interval between 1902 (the median year of historic samples used in this analysis) and 2014 (contemporary). The right-hand panel shows the selection coefficients under the balancing selection model that are consistent with observed changes in hb-chr2 inversion frequencies in Europe. Across the range of possible polymorphic equilibrium frequencies under balancing selection, equilibria near unity (*e.g*., *p_eq_* = 0.98) require modestly strong selection (average *S* = 0.028; 95% CI = [0.017, 0.043]) to explain the observed frequency changes in hb-chr2, and lower equilibrium states require stronger selection.

**Figure S18.**
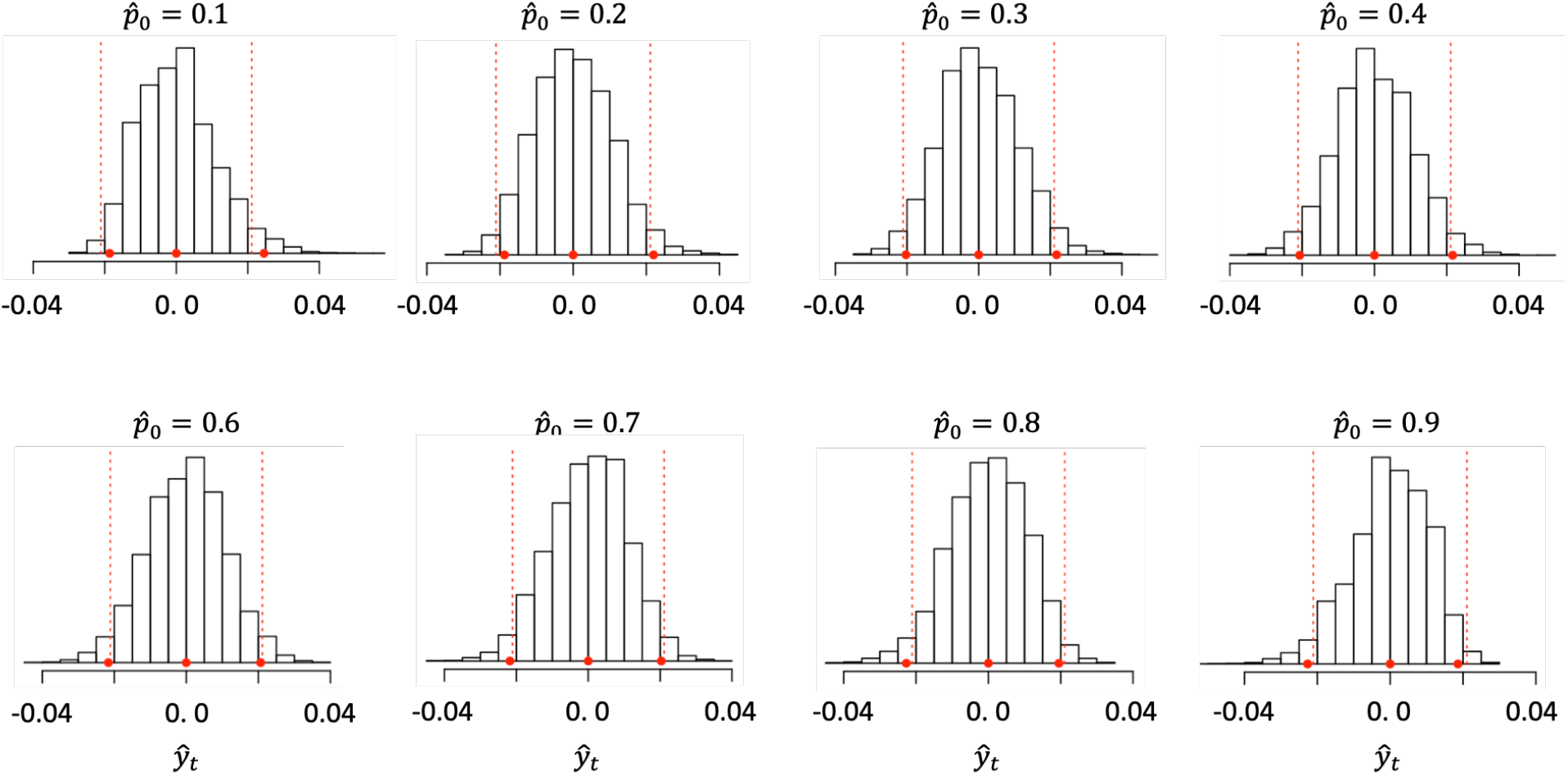
Simulated distribution of the scaled metric of divergence, 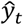, given different estimated values of the initial (historic) allele frequency. Each distribution is based on 10^4^ independent and neutrally evolving SNPs. The simulations use *n*_0_ = 182, *n_t_* = 312, and *t* = 131, with a moderate effective population size (*N_e_* = 10^4^). Histograms show the distribution of 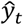 estimates, the red circles show the simulated mean and 95% confidence intervals for simulated data, and the vertical broken red lines show +/− 1.96 SD, where SD is the square root of our analytical expression for 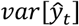.

## SUPPLEMENTARY TEXT

### S1: Estimating the strength of selection on hb-chr2 using temporal changes

The putative inversion hb-chr2 increases in frequency between historical and contemporary European populations, consistent with selection favouring its increase in the invasive range. To infer strengths of selection that are sufficient to explain the pattern of frequency change, we considered two simple, deterministic models of selection for the inversion:

- A *positive selection model* in which the inversion is favoured over the standard haplotype and is eventually expected to fix in European populations
- A *balancing selection model* in which the increased frequency of the inversion brings it closer to a hypothetical polymorphic equilibrium within the European range.

We note that, while genetic drift will inevitably play some role in the allele frequency dynamics of loci subject to selection (because populations are finite in size), evolutionary dynamics are well-approximated by deterministic models provided the allele frequencies of favourable variants are at least moderately common in the population and selection is strong relative to the inverse of the effective population size^81^. Both assumptions are easily met, and should apply to the hb-chr2 haplotype.

#### Positive selection model

Let *p* represent the frequency of the hb-chr2 inversion haplotype. Under a model of positive selection with no dominance, the general solution for the ratio of inversion to standard haplotype frequencies (the ratio defined as *x* = *p*/(1 – *p*)) is:

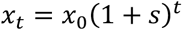

which is easily rearranged to solve for the frequency of the inversion:

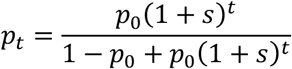

(*e.g*. ^82^ pp. 200-203), where *s* is the fitness increase associated with each copy of the inversion (*i.e*., the fitnesses of inversion heterozygotes and homozygotes, relative to individuals without the inversion, are 1 + *s* and (1 + *s*)^2^, respectively). Under this parameterization, the difference in relative fitness between inversion and standard haplotype homozygotes is 2*s*(1 + *s*/2) ~ 2*s*, with the 2*s* approximation applying when *s* is small (as we infer below).

The strength of selection is a function of the inversion frequency shift from *p*_0_ to *p_t_* following *t* generations of evolution:

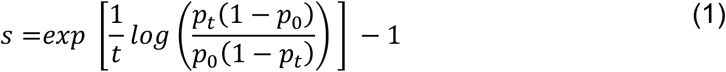

Since common ragweed is an annual plant, *t* refers to the number of years that have transpired, and *p*_0_ and *p_t_* can be estimated (with uncertainty) from the contemporary and historical samples.

We used eq. (1) to infer strengths of selection (*s*) that would be consistent with the estimated change in inversion frequencies over time, focusing on the estimated frequencies at the midpoint of the European range (table S20). Inversion frequencies were estimated as p = 0.37 (95% CI = {0.30, 0.45}) in the historic samples, with 1902 representing the median date of samples included in the analysis. For modern samples (sampling date of 2014), the estimated frequency was p = 0.69 (95% CI = {0.60, 0.76}). Given the large sizes of historic and modern collections, and the fact that haploblock frequencies remain intermediate across time, the distance between the 95% CI bounds will be roughly 3.92 standard errors of each frequency estimate. To take into account uncertainty in the frequency estimates, we simulated values of *p_t_* (modern) and *p*_0_ (historic) and used these values, along with eq. (1), to generate a distribution of selection coefficients (*s*) consistent with our data. Specifically, for a given time interval (i.e., historic or modern), we drew 10^6^ pseudo-random numbers from a normal distribution with mean corresponding to the point estimate of the haplotype frequency for the interval (i.e., 0.37 for historic; 0.69 for modern), and a standard deviation of *d*/3.92, where *d* is the difference between the 95% CI of the estimate. Our analysis yielded 10^6^ values of *s* that were compatible with the set of simulated inversion frequencies. The estimate and 95% confidence interval for *s* was calculated directly from the distribution of *s* values.

#### Balancing selection model

Although temporal changes in the hb-chr2 haplotype are consistent with changes predicted under the positive selection model presented above, we wished to also evaluate an alternative model in which balancing selection favours evolution of the inversion towards an equilibrium polymorphic state. To explore the strength of selection towards a hypothetical equilibrium within the European range, we considered a simple model of overdominant selection. Note that the overdominant selection model is dynamically equivalent to many other balancing selection models provided the differences in fitness among genotypes are small (consistent with our analysis below). Our results based on the overdominance model should, therefore, apply more broadly to other scenarios of balancing selection, including scenarios involving negative frequency-dependence and antagonistic pleiotropy^83,84^.

Following standard theory (*e.g*., ^85^ pp. 270-272), the expected change in frequency over a generation (generation *t* to generation *t* + 1) is:

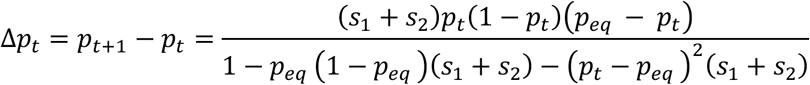

where *s*_1_ and *s*_2_ refer to the fitness costs of being homozygous for inversion and standard haplotypes, respectively, and *p_eq_* is the equilibrium frequency of the inversion. Using a continuous-time approximation, we can solve for the overall selection coefficient, *S* = *s*_1_ + *s*_2_, that is consistent with a frequency shift from *p*_0_ to *p_t_* < *p_eq_* across *t* generations:

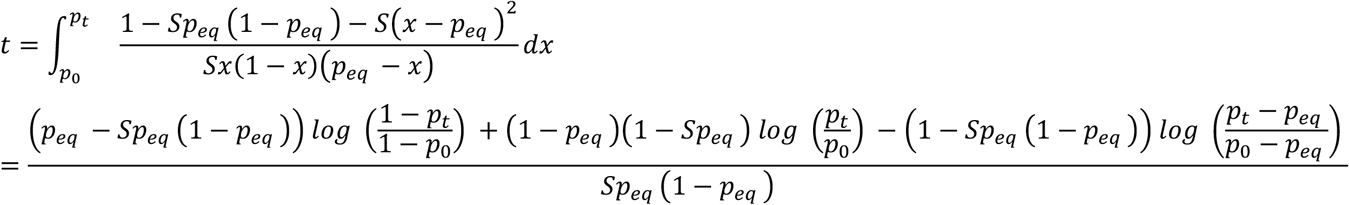

Solving for *S*, gives us:

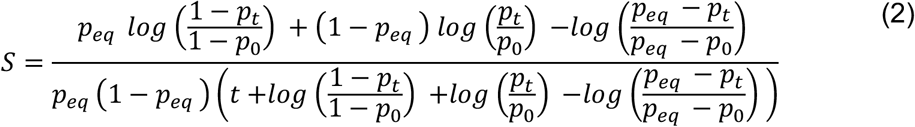

To infer the strength of selection (*S*) that would be consistent with observed inversion frequencies and a given equilibrium value (*p_eq_*), we simulated 10^6^ inversion frequencies consistent with the estimated frequency and its sample size at historical time point (~1902) and 10^6^ frequencies consistent with the estimate for the contemporary sample. (Frequencies were simulated as described in the positive selection model section, above). We used each pair of simulated inversion frequencies and eq. (2) to infer the value of *S* consistent with the frequency values. The resulting distribution of 10^6^ simulated *S* values was used to calculate 95% CI for *S* consistent with the data. We focused on equilibrium values outside of the 95% CI for contemporary inversion frequencies (*i.e*., values of *p_eq_* between 0.80 and 1). The results show that plausible selection coefficients under scenarios of balancing selection are consistently greater than those of the positive selection model (fig. S17). Selection under the positive selection model can, therefore, be regarded as a lower bound for the strength of selection consistent with the observed temporal changes in European hb-chr2 inversion frequencies.

### S2: Selection estimated from spatial changes in haploblock frequency

#### Cline theory

We will consider the simplest possible population genetics model of local adaptation in a species that is continuously distributed along a single axis of space (*e.g*., from north to south), with *x* representing location along the axis, and *x* = 0 representing a specific point in space where the environment relevant to selection at a focal locus—in this instance, a genomic region segregating for an inversion—changes abruptly. We assume that the inversion is favoured in locations where *x* > 0 (*e.g*., in the north) and the standard haplotype is favoured in locations where *x* < 0 (*e.g*., the south). We further assume that population density is uniform across the spatial gradient (at least within the vicinity of the environmental transition), and that individual dispersal follows a symmetric, Gaussian distribution with variance of *σ*^2^ (the unit of distance is arbitrary, though *σ* and *x* should have the same units, *e.g*.: if distance in *x* is measured in kilometres then *σ* should also be expressed in km; *σ*^2^ corresponds to the migration rate, *m*, between adjacent patches in discrete stepping stone models)^86^.

Given the stated assumptions, the inversion frequency dynamics at location *x* can be described using the following reaction diffusion equation:

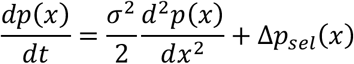

where Δ*p_sel_*(*x*) is the local response to selection^44,87^. With symmetrical strengths of selection at each side of the environmental transition, and no dominance, then Δ*p_sel_*(*x*) ≈ *sp*(*x*)(1 − *p*(*x*)) within the northern region of the range where the inversion is favoured, and Δ*p_sel_*(*x*) ≈ −*sp*(*x*)(1 − *p*(*x*)) in the southern portion of the range where the standard haplotype is favoured; both expressions are valid for modest-to-weak selection (0 < *s* < ~0.1). As in the positive selection model presented above, this parameterization leads to local fitness differences of ~2*s* between inversion and standard haplotypes. Incorporating dominance does not change our results provided the dominance relations between the alleles are consistent across the range (*i.e*., under “parallel dominance”^88^). At equilibrium between selection and migration, the maximum cline slope will be:

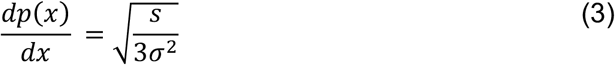

Following Roughgarden^87^, the equilibrium general solution for the cline is:

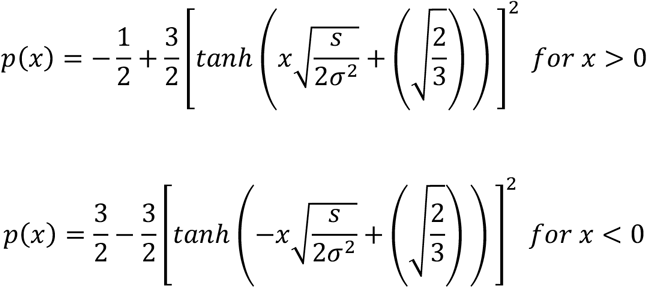

#### Estimating cline slopes by logistic regression

A logistic regression model for inversion frequency as a function of geographic location (*x*) is:

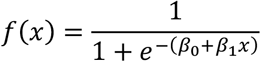

The parameters of the model (*β*_0_ and *β*_1_) can be estimated by fitting the data to the log-odds (logit):

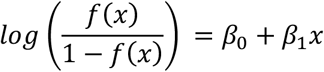

Using the theoretical cline functions (above) to calculate the log-odds, we obtain the following slopes. For shallow clines—those with a geographically broad clinal region, where the maximum slope can be accurately estimated—we have:

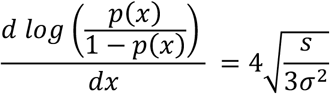

For steep clines—those with a narrow clinal region, where the maximum slope will be underestimated using the logit function—we have:

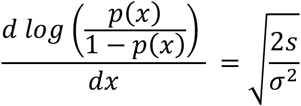

We get the following estimates from these two limits:

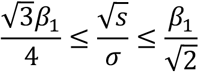

Given the point estimates and 95% CI for *β*_1_ in table S16, we can calculate plausible ranges for the lower bound of 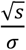 by multiplying the values for *β*_1_ by 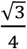

#### Comparisons of spatially varying selection among haploblocks

All of these estimates rely on the assumption that the system is at equilibrium within each range and time point, though that assumption may be more valid for some cases then others. To the extent that it is a reasonable assumption, and if effects of gene flow are consistent across the genome, we can estimate the relative strength of spatially varying selection on different haploblocks (e.g., haploblocks arbitrarily labeled “A” and “B”) as:

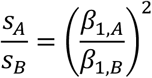

### S3: A simple null model of temporal allele frequency changes under drift

To evaluate whether temporal changes in candidate loci exceeded neutral expectations under drift in the absence of selection, we compared the distribution of the following standardized measure of divergence for a large sample of putatively neutral SNPs with the same metric calculated for selection candidates. Let divergence after *t* generations be defined as:

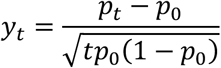

where *p*_0_ and *p_t_* represent the initial and final frequencies of an allele at a bi-allelic locus. We shall show below that, provided loci with low minor allele frequencies are first filtered out of the analysis, the metric follows a symmetric distribution that is approximately independent of the initial frequency.

For a locus with initial frequency of *p*_0_, the frequency after one generation of drift is given by:

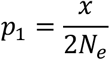

where *x* is a random variable drawn from a binomial distribution with parameters 2*N_e_* and *p*_0_, where *N_e_* is the effective population size (which follows the standard, Wright-Fisher model of genetic drift). The expected value and the variance for *p*_1_ is therefore *p*_0_ and *p*_0_(1 − *p*_0_)/2*N_e_*, respectively. The model can be extrapolated for a modest number of generations, after which the allele frequency (*p_t_* after *t* generations) has an expected value of *p*_0_ and variance of *tp*_0_(1 − *p*_0_)/2*N_e_*. The latter will eventually break down as *t* increases, but it should be appropriate provided *t*/2*N_e_* is small and the initial frequency is not too close to zero or one, as we assume below. From these expressions, the standardized measure of allele frequency divergence in the population under drift (and no selection) has an expectation of zero and a variance of:

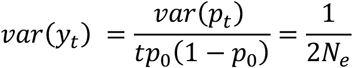

which is independent of the initial frequency.

In reality, error in the estimates of *p*_0_ and *p_t_* will also affect the test statistic, and this will tend to inflate the variance, but doesn’t alter the conclusion that (under a null model of drift with no selection) the distribution of the estimates of *y_t_* (which we denote as 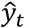) will be roughly independent of the initial allele frequencies in the historic sample. If we define 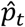 and 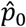 is the estimates of the allele frequencies, then our test statistic is:

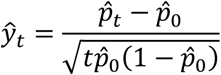

The mean and variance of 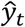 can be calculated using the following steps:

*Step 1*. The expected value and variance of 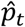 conditioned on the final population frequency (*i.e*., the true frequency, *p_t_*) is:

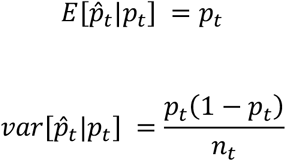

where *n_t_* represents the number of genes sampled in the contemporary population (*e.g*., for hb-chr2, 156 individuals were sampled for the contemporary estimate in Europe; given diploidy, we have *n_t_* = 312).

*Step 2*. The expected value and variance of 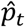 conditioned on the initial population frequency (*i.e*., the true frequency, *p*_0_) is:

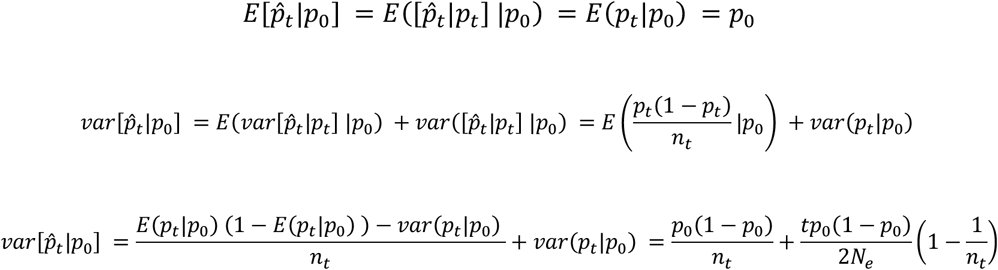

where *N_e_* is the effective population size, and *n*_0_ is the number of genes sampled in the historic population.

*Step 3*. Among loci with an initial frequency estimate of 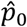, the true initial frequency (*p*_0_) will, roughly, follow a distribution with mean and variance of 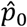 and 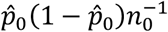, respectively. Consequently, the expected value and the variance of 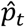 conditioned on the initial frequency estimate 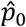 will be:

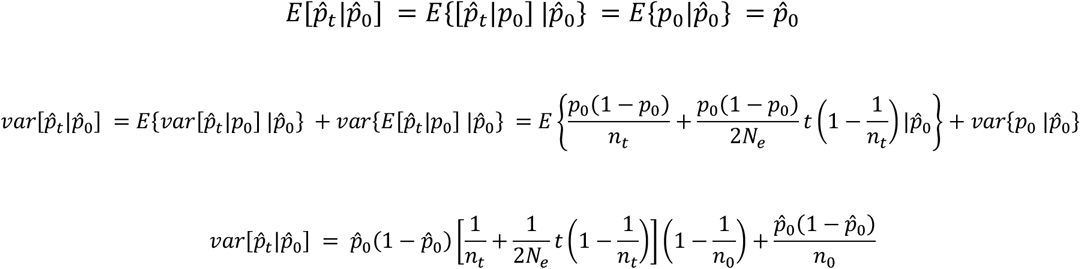

Therefore, the expected value and the variance for 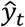, given an initial frequency estimate of 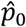, will be:

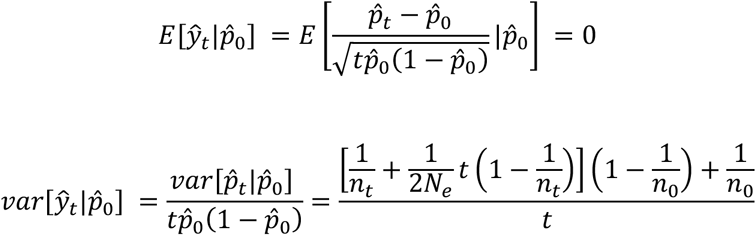

Note that the final expressions are, once again, independent of the initial frequency, though (once again) the pathway to these results requires that 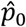 is not too close to zero or one. Because of this independence, we can pool loci with different initial frequency estimates (with pooling after loci with low minor allele frequency are first removed) to approximate the null distribution for 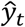 as well as the variance of the test statistic: 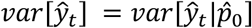.

Incidentally, the expression for 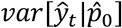 can be rearranged by solving for the effective population size across the *t* generations, *i.e*.:

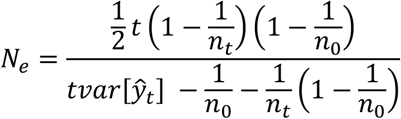

A rough estimate of *N_e_* can be obtained from a set of independent neutral SNPs by using the above formula with the estimated variance of 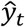 substituted for 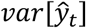.

#### Simulations

We carried out simulations to test the theoretical predictions of the neutral model presented above, and found that they work well as long as the initial allele frequency estimates are not too close to zero or one (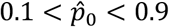 performs well and 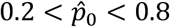 is excellent). Simulations for a given value of 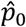 were carried out using the following steps. First, we used rejection sampling to simulate a distribution of initial population frequencies (*p*_0_) for a given value of 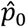. For each SNP, we sampled a true population frequency (*p*_0_) from a neutral stationary distribution (*i.e*., a single draw from a symmetric beta distribution with parameters theta = 0.05, which corresponds to the population-scaled mutation rate for the locus)^89^. We then generated a frequency estimate for the SNP from a single draw from a binomial distribution with parameters *p*_0_ and *n*_0_ = 182, where *n*_0_ is the number of genes sampled in the historic population. We retained the first 10^4^ simulated SNPs whose estimate after binomial sampling matched the focal value of 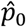. From the retained SNPs, we carried out forward Wright-Fisher simulations under pure drift for *t* generations to determine the contemporary population frequency (*p_t_*) for each SNP. We then carried out a second round of binomial sampling (with parameters *p_t_* and *n_t_*) for each SNP to generate a final allele frequency estimate. The frequency estimates were used to calculate 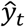 for each simulated SNP. Fig. S18 shows simulated distributions of 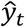 for different values initial frequency estimates 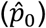. The distributions are roughly independent of 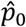 and their 95% CI are well-approximated by the 95% confidence interval predicted by a normal distribution with variance corresponding to our analytical expression for 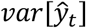 (see above).

